# Functional diversity in the Hsp60 of *Sulfolobus acidocaldarius*: mosaic of Group I and Group II chaperonin

**DOI:** 10.1101/2024.01.14.575554

**Authors:** Koustav Bhakta, Mousam Roy, Shirsha Samanta, Abhrajyoti Ghosh

## Abstract

External stress can disrupt protein homeostasis in organisms, necessitating the involvement of heat shock proteins (Hsps) to restore equilibrium and ensure survival. Unlike other organisms, the thermoacidophilic crenarchaeon *Sulfolobus acidocaldarius* lacks Hsp100, Hsp90, and Hsp70, possessing only two small heat shock proteins (Hsp14 and Hsp20) and one group II chaperonin, Hsp60. This raises questions about how protein substrates are protected and transferred to Hsp60 for refolding without other chaperones. Our study focused on ATP-dependent Hsp60 in *S. acidocaldarius*, revealing its formation of oligomeric structures in the presence of ATP. While ATP hydrolysis is not essential for oligomer formation and lid closure, it is crucial for Hsp60’s chaperone activity, effectively folding stress-denatured substrate proteins by stabilizing their folded conformations. The mechanism involves hydrophobic recognition of unfolded substrates, encapsulating and releasing them in a more folded state. Negatively charged inner surface of the ring seems to be responsible for driving the folding of the substrate. Importantly, Hsp14 was found to transfer substrate proteins to Hsp60αβ, orchestrating their refolding into an active state. Beyond protein folding, Hsp60β protects the membrane under stress, contributing to maintaining membrane rigidity. Hsp60 exhibits nested cooperativity in ATPase activity, adapting to ATP concentration changes and interestingly Hsp60β and Hsp60αβ complex shows a mosaic behaviour during ATP hydrolysis belonging to both group I and group II chaperonin respectively. In conclusion, our study provides insights into the intricate mechanisms employed by Hsp60 in *S. acidocaldarius* to maintain protein homeostasis. It offers a comprehensive understanding of Hsp60’s role in the heat shock response pathway, shedding light on fundamental cellular processes in extremophilic archaea.

## Introduction

Chaperonins are highly conserved protein complexes in all domains of life. They play a crucial role in the correct folding of proteins and act as guardians against the harmful aggregation of improperly folded or misfolded proteins [1–3]. Chaperonins are often up-regulated in response to cellular stress conditions, such as elevated temperatures, oxidative stress, nutrient stress, and are consequently categorized as heat-shock proteins. These specialized proteins help maintain proteostasis by assisting in protein folding and preventing the aggregation of misfolded proteins when cells are under stress. Chaperonins are composed of multiple copies of one or more subunits, which assemble into molecular cages with a mass of approximately 1 MDa. These cages serve as protective chambers, sequestering misfolded protein substrates within their inner cavities. The chaperonin function relies on allosteric binding and the hydrolysis of ATP, which are thought to induce ratchet-like movements within the cage structure [4–7]. These dynamic conformational changes aid in properly folding proteins entrapped within the chaperonin complex. These movements are believed to initiate alterations in the substrate-binding surfaces, facilitating the overcoming of local energy minima and promoting the correct substrate folding. Chaperonins can be classified into two groups, namely group I chaperonin and group II chaperonin. Group I chaperonins comprise 14 copies of a 60-kDa protein arranged with 72-point symmetry [8, 9]. These chaperonins are found in eubacteria, known as GroEL, and are also present within the mitochondrial matrix, referred to as cpn60/HSP60. The Escherichia coli chaperonin, GroEL, is a well-studied chaperonin that assists in protein folding. Working in conjunction with its cofactor, GroES, in an ATP-dependent process, GroEL acts as a folding chamber, and GroES serves as a lid to cap the cavity of GroEL. This coordinated action facilitates the release of substrate proteins, allowing them to undergo folding within the GroEL-GroES cavity [10]. On the contrary, group II chaperonins are found in archaea and eukaryotes. In archaea, they are popularly known as thermosomes (TF55), while in eukaryotes, they are known as CCT (chaperonin-containing TCP-1) or TRiC (T-complex protein-1 ring complex). Group II chaperonins, lacking a counterpart to GroES, exhibit unique features. The crystal structure of a Group II chaperonin from the acidothermophilic archaeon *Thermoplasma acidophilum* indicates that long helical protrusions from the apical domain serve as a “built-in-lid”, replacing the function of GroES and enclosing the central cavity [11, 12]. Certain archaeal chaperonins are assembled from a single type of 60-kDa subunit, while the majority are composed of two paralogous subunits, known as subunit α and subunit β [4, 12]. These subunits are arranged alternately in a hexadecameric complex, forming two eight-membered rings [13–16]. Importantly, these archaeal chaperonin subunits share a close similarity with the eight paralogous subunits found in eukaryotic CCT complexes, which are also organized into pairs of eight-membered rings [1, 3, 4, 17, 18]. So, it is evident that these group II chaperonins from eukaryotes and archaea share an evolutionary relationship [1, 3, 19].

Thermophilic Factor 55 (TF55), also known as rosettasome, is a group II chaperonin found in the hyperthermophilic archaea *Sulfolobus* [20]. *Sulfolobus* possesses three paralogous TF55 subunits, namely subunits α, β, and γ, which come together to create a variety of distinct complexes [21]. The β subunit of *Sulfolobus* was the first group II chaperonin subunit to be characterized, primarily due to its elevated expression levels in response to heat shock conditions [22]. The up-regulation of the β subunit in *Sulfolobus* has been identified as a critical factor enabling the organism to withstand extreme temperatures, including those as high as 92°C [20, 23]. Electron microscopy confirmed that these particles form a double-membered ring structure with 9-fold symmetry [24, 25]. However, when the α subunit was identified, it led to the conclusion that *Sulfolobus* TF55 is most likely assembled from α and β subunits, following a similar pattern observed in other archaeal chaperonins that exhibit 8-fold symmetry [26, 27]. More recently, the discovery of the γ subunit in the *Sulfolobus* genome [28] and following gene expression analysis of *Sulfolobus* cells revealed that the relative proportions of each of these subunits vary with temperature [26]. Specifically, the third subunit, γ, is only found under cold-shock conditions (60°C–75°C) whereas subunits α and β are present during normal growth temperatures (75°C–79°C). In contrast, the β subunit becomes the predominant subunit under heat-shock conditions (>86°C).

Our study explores the group II chaperonin of *Sulfolobus acidocaldarius*, a thermoacidophilic crenarchaeon thriving at 75-80°C and pH 2-3. In *Sulfolobus*, the heat shock protein machinery is minimal, consisting of two small heat shock proteins (Hsp14 and Hsp20) and one group II chaperonin, Hsp60, with three subunits (α, β, and γ). Notably, there is a temperature-dependent dynamicity in chaperonin complex formation. At 75 °C, α and β subunits form a hexadecameric complex [21]; at lower temperatures, α, β, and γ form a octadecameric complex [21]. However, at temperatures above 85 °C, the β subunit predominates, leading to the formation of octadecameric homo-oligomeric complex [21]. The ATP-independent small heat shock proteins lack substrate refolding ability, leaving this task to the only ATP-dependent group II chaperonin, Hsp60. We characterized the Hsp60β complex, finding that ATP binding induces oligomer formation and lid closure, while ATP hydrolysis being necessary for its chaperonin activity, protecting substrates from stress-induced aggregation and facilitating folding. Unfolded substrate recognition and folding involve hydrophobicity and charge-driven interactions respectively. Besides its chaperonin function, Hsp60β also protects membranes from stress-induced damage. The α subunit forms a hetero-oligomeric complex with β, protecting substrates from stress-induced aggregation and additionally it receives assistance from Hsp14 in unfolded substrate transfer. Finally, we assessed the ATPase activity of Hsp60β and Hsp60αβ complex and found out that there lies a nested cooperativity in their ATPase activity. Intriguingly, we observed that the ATPase activity of homo-oligomeric Hsp60β resembles that of group I chaperonin from bacteria while hetero-oligomeric Hsp60αβ resembles that of group II chaperonin from eukaryotes, highlighting the mosaic nature of archaeal cellular machineries. This mosaic nature is not limited to phylogenetic evolution but also extends to their cellular machinery, emphasizing the distinctiveness of this life domain.

## Results and discussions

### Hsp60β forms oligomeric structure in presence of ATP and oligomeric Hsp60β protects substrate against stress induced aggregation

Our investigation explored the impact of Hsp60β overexpression on *E. coli* cells. *E. coli* cells with an empty vector could not withstand 50 °C heat stress, while those overexpressing Hsp60β thrived for 120 minutes, suggesting that overexpressed Hsp60β protects against stress-induced damage (Fig 1a-b).

**Figure 1:**
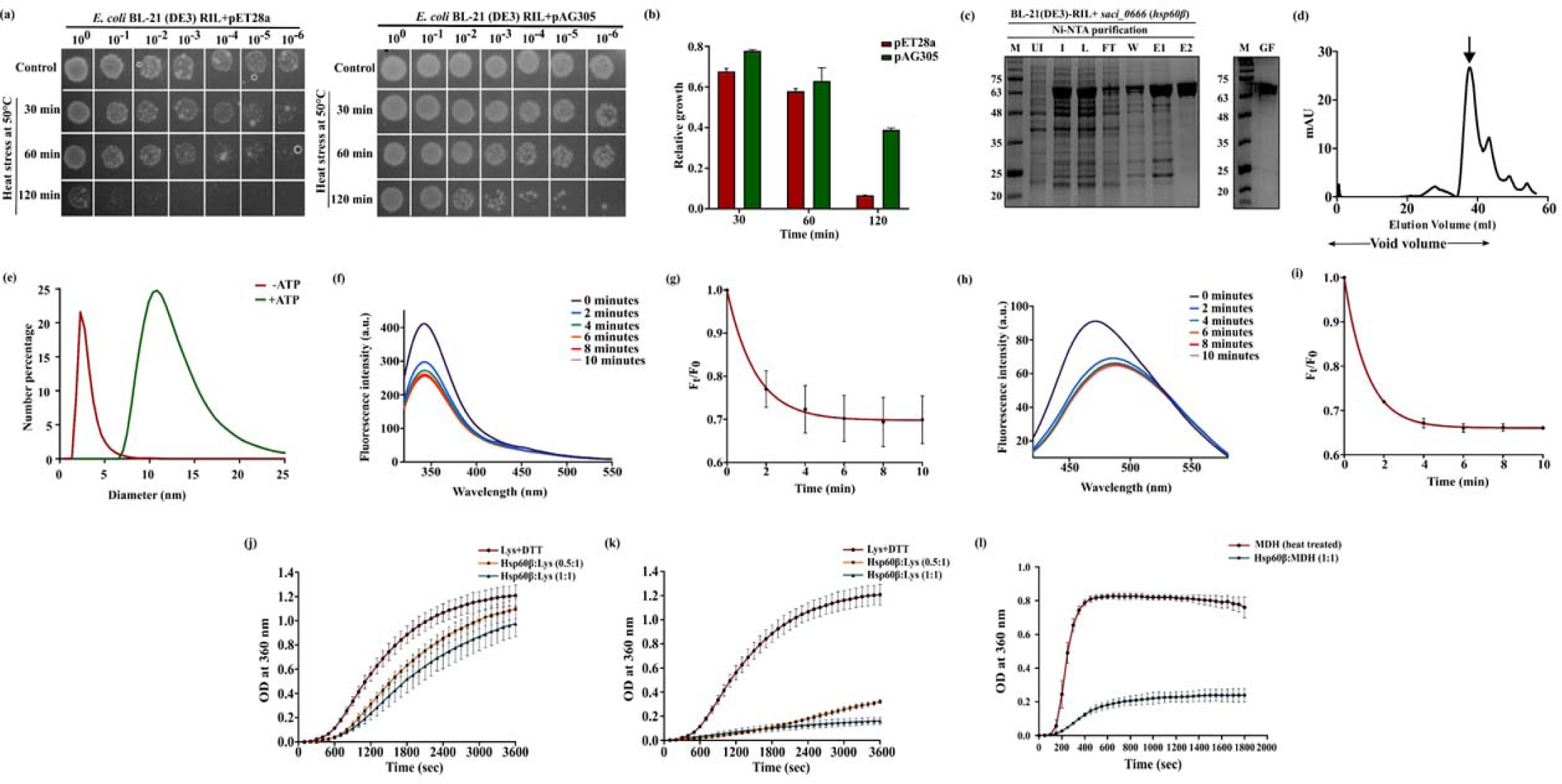
Overexpressed Hsp60β can protect *E. coli* from heat stress and it can form oligomeric structure in presence of ATP. (a) In this experiment, we expressed empty pET-28a vector and pAG-305 (containing *hsp60*β in pET-28a) in *E. coli* BL-21 (DE-3) cells and subjected them to a heat stress of 50 °C for up to 2 hours. Notably, the cells expressing the empty vectors succumbed to the stress, while those expressing Hsp60β remained viable throughout the entire duration of the test. (b) The spots were quantified with ImageJ software and relative growth was measured to quantitate the results. (c) Hsp60β was subsequently purified using affinity chromatography. The fractions are labelled as follows: M (marker), UI (uninduced), I (induced), L (load), FT (flow through), W (wash), E1 (elution 1), and E2 (elution 2). Following purification, the protein was subjected to gel filtration (GF). (d) Analysis of the gel filtration process indicated that the protein eluted in the void volume, suggesting the formation of oligomeric structures. (e) Dynamic Light Scattering (DLS) measurements revealed a significant shift in the hydrodynamic diameter in the presence of ATP, further supporting the formation of oligomers. (f) Examination of tryptophan fluorescence in Hsp60β indicates that in the presence of ATP, there is a progressive decrease in tryptophan fluorescence intensity with increasing time duration. (g) The rate constant for oligomer formation was derived from the decay curve of tryptophan fluorescence. (h) An analysis of ANS fluorescence reveals that over time, the ANS fluorescence intensity decreases, accompanied by a red shift in the spectrum, indicating the formation of oligomers with fewer exposed hydrophobic patches. (i) ANS fluorescence demonstrated a similar rate of oligomer formation obtained from tryptophan fluorescence. (j) In its monomeric form, Hsp60β is unable to shield lysozyme from aggregation induced by DTT at concentration ratios of 0.5:1 or 1:1. However, oligomeric Hsp60β can effectively protect both lysozyme (k) and MDH (l) from aggregation triggered by DTT and heat, respectively. Error bar represents SEM obtained from three or more sets of replicates.

Previous studies indicated that Hsp60/chaperonin formed oligomeric structures in the presence of ATP, with ATP-induced conformational changes in eukaryotic CCT and archaeal thermosome [21, 29–31]. We aimed to investigate Hsp60β’s potential to form oligomers under specific conditions. We treated purified Hsp60β (Fig 1c) with 5 mM ATP and 25 mM MgCl_2_, subjecting it to 80°C heat treatment for 10 minutes, a known oligomerization protocol [21]. Gel filtration revealed protein elution at approximately 40 ml (Fig 1c-d), affirming Hsp60β’s oligomerization in the presence of ATP and MgCl_2_. Dynamic light scattering showed Hsp60β with a hydrodynamic diameter of 2.5 nm without ATP, signifying a monomeric state. In contrast, ATP induced a shift to 12 nm, confirming oligomer formation (Fig 1e). Similar studies have been done in the hyperthermophilic archaeon *Sulfolobus shibatae*. Both α and β subunits of the chaperonin formed homooligomeric structures individually in the presence of ATP [26]. Chaston *et al*. further supported Hsp60β’s homo-oligomer formation, reporting an octadecameric chaperonin complex in *S. solfataricus* [21]. This variation in thermosome composition had implications for filament formation, suggesting its importance in archaeal structure and function [26]. It is suggested that the chaperonin filaments serve as a mechanism for sequestering double rings involved in protein folding. Alternatively, it is hypothesized that these filaments contribute directly to the structural integrity of cells [32].

Intrinsic tryptophan fluorescence studies were conducted to explore conformational changes associated with oligomerization. Tryptophan residues exhibited a consistent reduction in fluorescence intensity over time, indicating conformational changes linked to oligomer formation (Fig 1f). Homology modeling revealed that all tryptophan residues in the protein’s quaternary structure were on the surface and conformational change due to oligomer formation probably made it much more solvent-exposed. The ^1^L_a_ state of tryptophan is solvent-sensitive, favoring its dominance in emission and explaining the reduced emission intensity. Through a photoelectron transfer mechanism, tryptophan fluorescence is also quenched in proteins by primary chain amides and amino acid side chains, with tryptophan acting as the electron donor [33]. Collectively, these factors, alone or combined, contribute to the observed quenching of tryptophan fluorescence in our experiment. This phenomenon may stem from compaction of protein structure induced by the oligomer formation. The fluorescence decay data were fitted to a first-order decay equation, yielding a rate constant of oligomer formation to be 0.70±0.04 min^-1^ (Fig 1g). This rate constant provides insights into the kinetics of the conformational changes driven by oligomer formation.

Further insights into structural alterations were gained by investigating surface hydrophobicity using ANS. There was a decrease in fluorescence intensity along with a red shift in the ANS fluorescence spectra (Fig 1h), indicating a reduction in the exposure of hydrophobic regions on the protein’s surface and alterations in the polarity of the microenvironment surrounding the ANS-bound regions. These observations collectively suggest that over time, there is a decrease in the surface hydrophobicity of Hsp60β. We speculate that oligomerization, induced by the presence of ATP, appears to drive these changes in hydrophobicity, potentially through the reorganization of the protein’s structure and surface properties. The rate constant for this conformational change due to oligomer formation was determined to be 0.77±0.01 min^-1^ (Fig 1i). The rates of oligomer formation, as determined by both tryptophan fluorescence and ANS fluorescence, are close to each other.

Research has demonstrated that *E. coli* GroEL can effectively inhibit the aggregation of various model polypeptides when transferred from a denaturing environment into a refolding buffer [34, 35]. Our investigation into the chaperone activity of the Hsp60β subunit centered on its ability to prevent protein aggregation, focusing on lysozyme and malate dehydrogenase (MDH). In the presence of DTT, lysozyme’s disulfide bonds are reduced, causing it to lose its native structure and form large, aggregated structures in the solution [36, 37]. Our findings revealed that Hsp60β, in the absence of ATP, did not exhibit the ability to protect lysozyme from DTT-induced aggregation (Fig 1j). However, adding ATP with Hsp60β significantly reduced lysozyme aggregation, with protection directly proportional to Hsp60β concentration (Fig 1k). This highlights Hsp60β’s role, along with ATP, in preventing chemical-induced lysozyme aggregation.

Elevated temperatures induce substrate protein unfolding and aggregation through hydrophobic interactions, forming particulates. MDH, prone to aggregation, exhibited reduced aggregation in the presence of ATP bound Hsp60β in a 1:1 ratio (at 50 °C). Hsp60β’s chaperone function probably preserved MDH’s structural integrity, preventing aggregation at high temperatures, as indicated by reduced scattering intensity (Fig 1l).

Our findings align with observations of chaperonins from different archaea. Previously, *Methanococcus maripaludis* chaperonin (Mm-Cpn) was shown to refold human γD crystallin, preventing aggregation induced by denaturing agents [38]. Similarly, *Methanosarcina mazeii* chaperonin (MmThs) shielded luciferase and rhodanese from aggregation induced by guanidinium chloride [39]. These parallel findings across different archaeal organisms highlight the evolutionary conservation of chaperonin-mediated protein protection mechanisms and underscore the importance of these molecular chaperones in safeguarding protein structure and function under challenging environmental conditions.

### ATP binding is sufficient for oligomer formation but ATP hydrolysis in necessary for its chaperone activity

Group II chaperonins are complex, ring-shaped structures dependent on ATP that interact with non-native polypeptides, aiding in protein folding across archaea and eukaryotes. They incorporate a built-in lid, which envelops substrate proteins within the central chaperonin chamber [40]. In our study, we observed that in presence of ATP Hsp60β formed oligomeric structure. However, our goal was to investigate whether ATP binding alone is sufficient for Hsp60β’s oligomer formation or if ATP hydrolysis is essential. To address this query, we conducted experiments to assess the oligomerization of Hsp60β in the presence of AMP-PNP, a “nonhydrolyzable” nucleotide analog that mimics the ATP ground state, possessing tetrahedral geometry like the slowly hydrolyzable nucleotide analog ATP-γ-S [41].

DLS data suggested that Hsp60β formed an oligomer with AMP-PNP, evident from a shift in hydrodynamic diameter (Fig 2a). Like ATP, oligomer formation by AMP-PNP induced conformational changes, reducing tryptophan fluorescence intensity (Fig 2b) and the rate of oligomer formation was measured to be 0.80±0.035 min^-1^ (Fig 2c). ANS fluorescence spectra corroborated these observations, indicating reduced hydrophobic exposure (Fig 2d). The rate of oligomer formation measured from ANS fluorescence was calculated to be 0.85±0.049 min^-1^ (Fig 2e). The rates of oligomer formation observed in this context closely resemble those achieved in the presence of ATP. These results demonstrated that AMP-PNP binding was sufficient to induce oligomerization and conformational change of Hsp60β. Thus, we can infer that ATP binding alone is adequate for forming oligomeric structures by Hsp60β and that ATP hydrolysis is not necessary.

**Figure 2:**
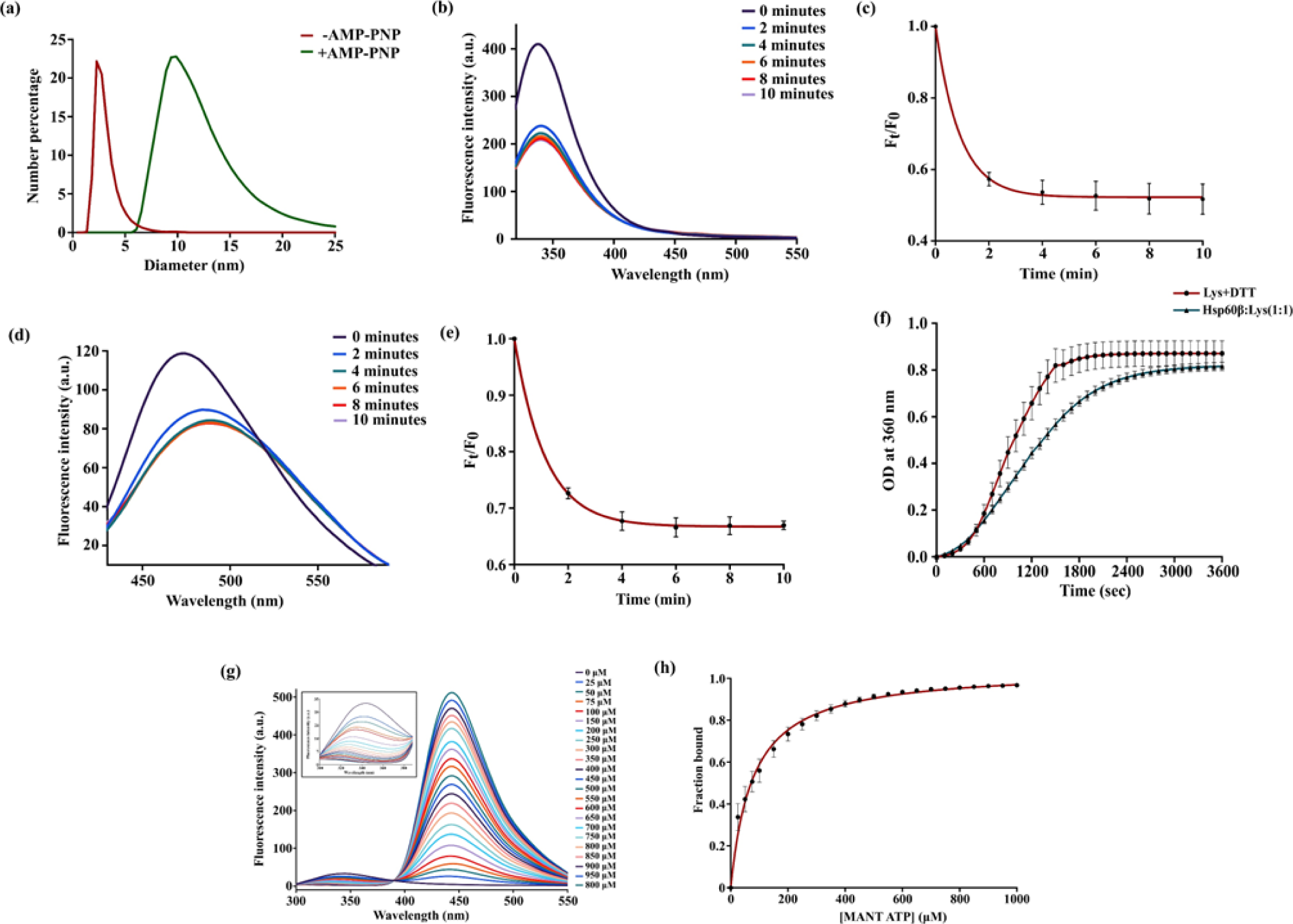
Only binding of ATP is sufficient for oligomer formation but ATP hydrolysis is necessary for its chaperone activity. (a) Data obtained from Dynamic Light Scattering (DLS) demonstrates that in the presence of the non-hydrolysable ATP analogue AMP-PNP, Hsp60β can indeed form oligomeric structures. (b) Tryptophan fluorescence measurements of Hsp60β reveal that in the presence of AMP-PNP, there is a progressive decrease in tryptophan fluorescence intensity over time. (c) The rate constant for oligomer formation was determined from the decay curve of tryptophan fluorescence. (d) ANS fluorescence analysis illustrates that with an increasing duration of exposure to AMP-PNP, the ANS fluorescence intensity diminishes along with a red shift, indicating the formation of oligomers with fewer surface-exposed hydrophobic patches. (e) ANS fluorescence, demonstrated a similar rate of oligomer formation that obtained from tryptophan fluorescence. (f) The oligomers of Hsp60β formed in the presence of AMP-PNP are unable to protect lysozyme from aggregation induced by DTT. (g) Hsp60β was subjected to titration with increasing concentrations of MANT-ATP, resulting in FRET between MANT-ATP (the acceptor) and tryptophan (the donor). As a consequence, the fluorescence intensity of tryptophan gradually decreased, while the intensity of MANT fluorescence increased. (inset) The fluorescence spectra of tryptophan exhibited a decrease in intensity along with a blue shift (h) The binding of MANT-ATP to Hsp60β was quantified using equation 9 as described in the materials and methods section, revealing a dissociation constant (K_d_) of 77.2±3.8 µM. Error bar represents SEM obtained from three or more sets of replicates.

Interestingly, we observed that in the presence of AMP-PNP, Hsp60β could not protect lysozyme from DTT-induced aggregation, suggesting that ATP hydrolysis is absolutely necessary for protecting a substrate against aggregation and probably for refolding purpose (Fig 2f). While ATP binding alone induced oligomerization and conformational change, our data supports that ATP hydrolysis is essential for Hsp60β’s chaperone function, particularly in protecting protein substrates from aggregation. We also tested the ATP hydrolysis properties of Hsp60β in absence and presence of substrate and observed that ATP hydrolysis increases significantly in presence of substrate (supplementary figure S1). ATP hydrolysis probably releases the substrate in the inner cavity of the chaperonin leading to refolding and activation [40].

We wanted to study the binding affinity of ATP to Hsp60β by using Förster resonance energy transfer (FRET) which is a widely used technique in molecular biology to investigate molecular interactions and spatial changes within biological molecules. In this particular study, FRET was employed to probe the binding affinity of ATP with Hsp60β, using the fluorescent ATP analogue known as 2’/3’-(N-methyl-anthraniloyl)-adenosine-5’-triphosphate (MANT–ATP) and tryptophan residues as key components. MANT–ATP is a modified form of ATP, where a fluorescent group (MANT) is attached to the ribose part of ATP. This modification makes MANT-ATP capable of emitting fluorescence when excited by specific wavelengths of light. Importantly, the absorption spectrum of MANT-ATP peaks at 355 nm, which coincidentally overlaps with the emission spectrum of tryptophan residues found in Hsp60β, a protein of interest in this study. However, MANT-ATP’s own fluorescence emission peaks at 448 nm, which does not overlap with tryptophan’s emission. To assess how ATP binding affects the spatial arrangement of tryptophan residues in Hsp60β, a titration experiment was conducted. Hsp60β was exposed to increasing concentrations (ranging from 0 to 1 mM) of MANT-ATP (Fig 2g) and energy transfer between tryptophan and MANT-ATP was measured during this process. The presence of MANT-ATP had a significant impact on the fluorescence characteristics of tryptophan residues in Hsp60β. As MANT-ATP bound to the Hsp60β subunit, there was a dose-dependent reduction in the fluorescence intensity of the tryptophan residues along with a blue shift (Fig 2g and 2g inset). Simultaneously, there was an increase in the fluorescence emission from MANT-ATP itself. The data obtained from these experiments allowed the determination of the binding affinity (K_d_) between MANT-ATP and the Hsp60β subunit. In this case, the K_d_ was found to be approximately 77.2±3.8 µM, indicating a relatively strong binding interaction between MANT-ATP and Hsp60β (Fig 2h).

ATP plays a pivotal role in orchestrating the conformational transitions of group II chaperonins. This conformational transition occurs due to inward movement of the apical domain or the lid of the chaperonin, shifting the structure from an open-lid, substrate-binding state to a closed-lid configuration [30, 42]. This conformational change encapsulates an unfolded protein within the central cavity while inducing a conformational shift within that cavity. The necessity of ATP hydrolysis for this conformational change has been a subject of debate. [30, 43, 44]. Based on our results, we hypothesize that ATP hydrolysis may not be essential for the conformational change; instead, the binding of ATP alone might be sufficient for oligomerization and potentially lid closure. Supporting studies, such as those on *Thermococcus* strain KS-1’s group II chaperonin, reveal that both ATP and AMP-PNP induce the transition from lid-open to lid-closed forms [44]. The presence of either nucleotide protects the chaperonin’s lid from proteolysis, indicating efficient lid closure [44]. They also tested the conformational change through tryptophan fluorescence [44]. Similar observations are reported in case of group II chaperonin from *Thermoplasma acidophilum* [45] and yeast CCT [42], where either ATP or AMP-PNP binding shifts the conformation from open-lid to closed-lid. Small angle neutron scattering and Cryo-EM studies revealed such information [42, 45]. Additionally, another group’s findings propose a two-step folding process featuring an initial conformational shift upon cavity closure, followed by a final folding step [46]. Their results indicate that ATP binding initiates the initial closure of the cavity. However, conflicting reports suggest that ATP hydrolysis is crucial for lid closure and substrate release, essential for productive protein folding. This viewpoint argues that ATP binding induces a conformational change and domain movement but is insufficient for CCT lid closure; ATP hydrolysis is deemed necessary for achieving the closed-lid conformation [40].

### Oligomeric Hsp60**β** stabilizes the folded state of a substrate

After finding out the role of ATP hydrolysis in Hsp60β’s activity, our investigation focused on understanding the mechanisms behind its function, particularly in mitigating thermal stress on malate dehydrogenase (MDH). Subjecting MDH to heat-induced denaturation, we monitored light scattering at 360 nm to quantify aggregation under varying temperatures. In the presence of Hsp60β, a significant decrease in scattering intensity was observed, indicating its protective effect (Fig 3a).

**Figure 3:**
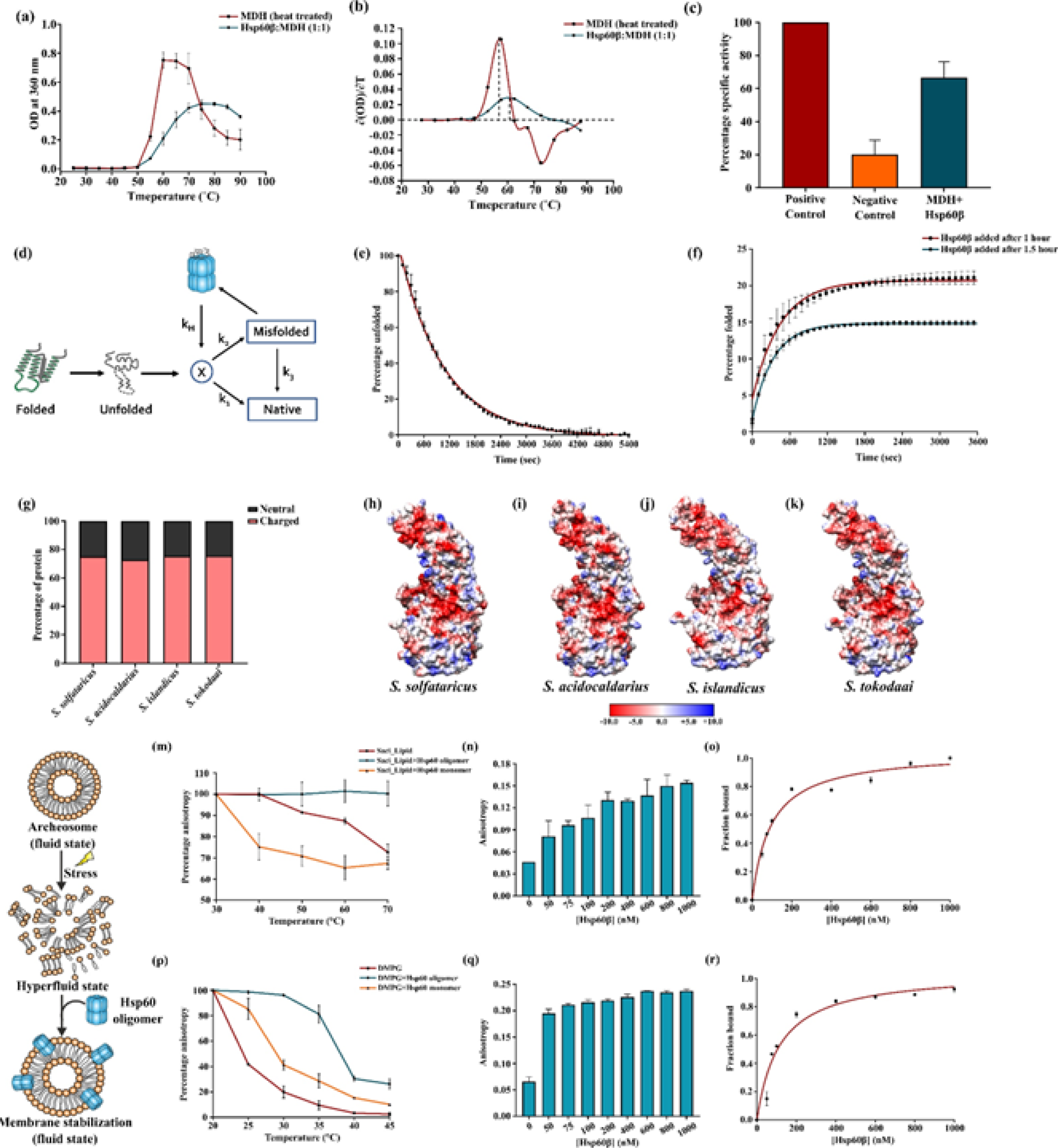
Hsp60β protects and stabilizes folded form of the substrate in an ATP dependent manner and recognizes substrate through hydrophobic interactions. It also stabilizes membrane. (a) Examination of thermal aggregation of MDH indicates that in the presence of Hsp60β oligomers, the scattering intensity decreases, which is indicative of protection against heat-induced damage. (b) Analysis of the first derivative plot of the scattering data reveals that in the presence of Hsp60β, the melting temperature (T_m_) of MDH increases, implying that Hsp60β stabilizes the substrate’s folded state. (c) The restoration of specific activity in MDH following heat treatment was assessed, and it was observed that in the presence of Hsp60β, the specific activity was restored up to 60%. (d) When a folded protein is subjected to stress, it undergoes a conversion into an unfolded state, which possesses high energy. Subsequently, these unfolded proteins rapidly adopt some secondary structure, transitioning to a lower energy state denoted as state X. From this state, the protein can either progress to a misfolded state (with rate constant k_2_) or return to its native state (with rate constant k_1_). The misfolded state can also revert to the native state with a rate constant of k_3_. Additionally, the misfolded protein has the capability to interact with a chaperonin, which can release the substrate back to state X with a rate constant of k_H_. This mechanism is referred to as the Iterative Annealing Mechanism (IAM). (e) Lysozyme was unfolded with DTT for 1.5 hours, and the unfolding process was monitored using ANS in a spectrofluorometer. The data was plotted as a percentage of lysozyme unfolded versus time. (f) Hsp60β was introduced after 1 hour and 1.5 hours. When Hsp60β was added after 1 hour, lysozyme was refolded up to 20% with a rate constant of 0.134±0.002 min-1. In the case where Hsp60β was added after 1.5 hours, lysozyme was refolded up to 15% with a rate constant of 0.179±0.003 min-1. (g) Analysis of the distribution of charged and neutral proteins in the proteomes of *S. solfataricus*, *S. acidocaldarius*, *S. islandicus*, and *S. tokodaai* reveals that in all these organisms, more than 70% of the proteins carry a net electric charge. Examination of the surface charge distribution of Hsp60β from (h) *S. solfataricus*, (i) *S. acidocaldarius*, (j) *S. islandicus*, and (k) *S. tokodaai* is presented. The crystal structure is available for Hsp60β from *S. solfataricus*, whereas the structures for the other organisms are homology-modelled structures. (l) The schematic illustrates the mechanism of membrane protection, where archaeosomes or DMPG LUVs are in a fluid state under physiological conditions. However, exposure to thermal stress triggers a transition into a hyper-fluid state. Hsp60β oligomers can interact with the membrane, preventing it from entering the hyper-fluid state and maintaining its stability. (m) and (p) As temperature increases, the percentage anisotropy of DPH associated with archaeosomes or DMPG decreases, indicating membrane fluidization. In the presence of Hsp60β monomers, the anisotropy also decreases, indicating that Hsp60β monomers do not provide protection. However, in the presence of Hsp60β oligomers, the percentage anisotropy of DPH does not decrease drastically, suggesting that Hsp60β incorporates into the membrane thereby maintaining its integrity. (n) and (q) Archaeosomes or DMPG LUVs were titrated with increasing concentrations of Hsp60β, and the raw anisotropy data was plotted. This demonstrates that as the concentration of Hsp60β increases, the anisotropy of DPH also increases, indicating the membrane incorporation of Hsp60β oligomers. (o) and (r) The titration data was fitted to a hyperbolic binding equation, revealing binding dissociation constants (K_d_) of 94.64±0.0025 nM and 119.33±6.22 nM for Hsp60β to archaeosomes and DMPG, respectively. Error bar represents SEM obtained from three or more sets of replicates.

To gain mechanistic insights, we calculated the melting temperature (T_m_) representing the inflection point between folded and unfolded states. Analyzing the first derivative of the thermal aggregation curve allowed us to pinpoint the T_m_. Under standard conditions, MDH exhibited a T_m_ of 56 °C±0.02, indicating susceptibility to thermal denaturation. In the presence of Hsp60β, the T_m_ increased to 60 °C±0.04, highlighting enhanced stability (Fig 3b). Moreover, Hsp60β restored 60% activity to heat-denatured MDH, demonstrating its ability to refold proteins (Fig 3c).

Together, these findings suggest that Hsp60β stabilizes the folded, functional state of MDH, preventing thermal-induced aggregation and denaturation. As a molecular guardian, Hsp60β protects the structural integrity and enzymatic activity of MDH at higher temperatures. This further highlights Hsp60β’s role as a crucial chaperonin in stabilizing protein structures and enhancing resilience to thermal stress, advancing our understanding of chaperone-client interactions in cellular protein quality control.

### Hsp60**β** recognizes substrate through hydrophobic interactions and folds substrate through charge driven interactions

We observed that Hsp60β plays a crucial role in stabilizing the partially folded state of the substrate protein. However, it is imperative to understand the recognition mechanism employed by Hsp60β to identify the misfolded state within the substrate protein. When a protein in its folded conformation encounters stressful or unfavourable conditions, it may undergo a transition to an unfolded state, typically characterized by higher energy levels. In this unfolded state, the protein rapidly assumes specific secondary structures, progressing towards a lower energy state referred to as “state X” (Fig 3d) [47]. From this state X, the protein has two potential pathways it can follow: it can either refold back into its native state, characterized by a rate constant denoted as “k_1_,” (Fig 2h) or it can misfold into an alternative, non-functional state, governed by a rate constant “k_2_” [47] (Fig 3d). It is worth noting that misfolded proteins also can revert to their native state, albeit at a slower rate, represented by the rate constant “k_3_.” Notably, misfolded proteins can interact with chaperonins, such as Hsp60, and these chaperonins can facilitate the return of the protein to state X, effectively initiating the folding process anew [47]. The release of the protein from the chaperonin back to state X is controlled by a rate constant denoted as “k_H_” (Fig 3d) and this step can happen repeatedly until the unfolded protein goes back to its native state from state X. This entire iterative process is known as the “iterative annealing mechanism” (IAM) [47].

A noteworthy aspect of the IAM is that it encompasses random, non-specific interactions between the chaperonin and the protein, regardless of its secondary structure [47]. Essentially, it postulates that these random, non-specific interactions between the chaperonin and the substrate protein are primarily driven by the exposure of hydrophobic residues [47]. The chaperonin actively seeks out these hydrophobic patches on the protein’s surface and binds to them, thereby assisting in the folding process.

To understand the recognition mechanism of Hsp60β, we investigated the chemical-induced unfolding of lysozyme and its refolding in the presence of Hsp60β (Fig 3e-f). The denaturation of lysozyme exposes hydrophobic patches on the protein’s surface. To quantify the increase in hydrophobicity, we employed the fluorescent probe ANS, which specifically binds to hydrophobic residues on the protein, increasing fluorescence intensity. We allowed lysozyme to unfold in the presence of DTT for one and a half hours. We then monitored and plotted the percentage of unfolded lysozyme over time (Fig 3e). To assess lysozyme refolding, we introduced Hsp60β at two distinct time points: after one hour and after one and a half hours of the unfolding process. When Hsp60β was added after one hour, we observed that lysozyme could refold up to 21% with a rate constant of 0.134±0.002 min^-1^ (Fig 3f). Similarly, when Hsp60β was introduced after one and a half hours, lysozyme exhibited a refolding percentage of up to 15% with a rate constant of 0.179±0.003 min^-1^ (Fig 3f). These findings suggest that adding Hsp60β later in time resulted in a slightly reduced effect on the maximum percentage of folded lysozyme. However, it is noteworthy that Hsp60β demonstrated a rapid recognition of the protein substrate revealed from the increased rate constant. This observation can be rationalized by the fact that Hsp60β recognizes hydrophobic patches on denatured protein substrates. The lysozyme subjected to unfolding for a duration of one and a half hours exhibited a higher ANS fluorescence intensity compared to the lysozyme unfolded for only one hour. So, it can be inferred that, the lysozyme that had been unfolded for one and a half hours presented more exposed hydrophobic patches than the one unfolded for just one hour. This increased exposure of hydrophobic regions likely contributed to the accelerated recognition of the substrate by Hsp60β. These results suggest that Hsp60β captures lysozyme through a non-specific general mechanism. We believe that hydrophobic interactions arising from differences in hydrophobicity between the unfolded and folded protein substrate play a pivotal role in this recognition process.

While hydrophobicity plays a significant role in the recognition of substrate proteins, our inquiry sought to uncover other potential interactions responsible for folding. To delve deeper, we analyzed the available proteome database of *S. solfataricus*, *S. acidocaldarius*, *S. islandicus*, and *S. tokadaai* on a comparative basis. Surprisingly, we discovered that the proteome of these members of the *Sulfolobaceae* family is characterized by a high degree of charge heterogeneity. Specifically, more than 70% of the proteins were found to be charged, while less than 30% were neutral (Fig 3g). This abundance of charged proteins may play a crucial role in intrinsic buffering within the cytosol, allowing maintenance of the internal pH of *Sulfolobus* (pH∼6.5) [48, 49] thriving in acidic environments (pH 2-3). However, previous studies have indicated that charged proteins, particularly those with a high net charge, are more susceptible to damage, mainly oxidative damage [50]. In the natural habitats of *Sulfolobus* species, oxidative damage induced by reactive oxygen species (ROS) is typical [51]. Such damage can alter the charge distribution of side chains, resulting in a loss of protein stability. Highly charged proteins are at a heightened risk of experiencing significant stability loss due to the compaction of charges into a smaller space during the folding process [50]. Oxidation of such highly charged proteins due to oxidative stress can lead to an overall negative charge and unfolding of proteins [50].

Interestingly, the Hsp60β subunit appears to have evolved to possess a patch of highly negatively charged regions on its internal surface. We scrutinized the crystal structure of *S. solfataricus* Hsp60β [21] and homology-modelled the structures of Hsp60β from *S. acidocaldarius*, *S. islandicus*, and *S. tokadaai* (Fig 3h-k). Remarkably, we observed the presence of a conserved negatively charged region within the inner surface of Hsp60β across all these structures, suggesting an evolutionary conservation of this feature. When a negatively charged unfolded protein enters the cage formed by Hsp60β, it encounters a repulsive force from the negatively charged wall of Hsp60β’s interior. This results in the compaction of the protein and the formation of a partially folded state, which is subsequently released into the surrounding environment. This protein can then undergo folding independently, or the iterative annealing mechanism can repeat itself. We hypothesize that the interplay between hydrophobicity-driven recognition and charge-driven folding of substrate proteins by Hsp60β works in tandem to complete the iterative annealing mechanism. This dual mechanism appears logical as it enhances the efficiency of substrate recognition and the subsequent protein folding process. Nevertheless, a dilemma arises in this mechanism when proteins expose positively charged residues upon unfolding; in such cases, hydrophobicity-driven recognition and compaction may constitute a significant driving force, but a thorough investigation is necessary to delve into the details.

### Hsp60**β** interacts and stabilizes archaeal and eukaryotic membrane lipids

In eukaryotic cells, Hsp60 is present in both the cytosol and mitochondria. During *Listeria monocytogenes* infection, mitochondrial Hsp60 is overproduced and associated with the plasma membrane [52]. In archaea, proteomic analysis of *Sulfolobales*’ secreted vesicles indicated the presence of Hsp60β, suggesting a role beyond protein stabilization, potentially affecting membrane lipids [53]. Given our prior observation that the expression of Hsp60β increases with rising temperature [23], we explored Hsp60β’s ability to associate with membrane lipids. To explore Hsp60β’s interaction with membranes (Fig 3l), we conducted experiments using *Sulfolobus* archaeosomes and DMPG (1,2-dimyristoyl-sn-glycero-3-phosphoglycerol) vesicles mimicking eukaryotic membranes. Given that Hsp60 belongs to the group II chaperonin class, it is logical to explore its interactions with archaeal and eukaryotic membrane lipids. Employing the fluorescent probe DPH, we measured steady-state fluorescence anisotropy, revealing insights into Hsp60β’s ability to interact with archaeal and eukaryotic membrane lipids.

In the absence of Hsp60β, archaeosome membrane fluidity increased with rising temperature, with a concomitant decrease in DPH anisotropy, reaching approximately 75% of its initial value at 70°C. However, when treated with oligomeric Hsp60β (ATP-bound), the membrane fluidization was significantly reduced, indicating a protective role of Hsp60β in stabilizing the archaeosome membrane. However, monomeric Hsp60β (without ATP) lacked this protective effect (Fig 3m). The association between Hsp60β and membranes was further confirmed by titrating archaeosomes with increasing Hsp60β concentrations, where we observed an increase in DPH anisotropy (Fig 3n). The association between Hsp60β and archaeosome was quantified with a hyperbolic binding equation, revealing a dissociation constant (K_d_) of 94.64±0.0025 nM (Fig 3o). These findings highlight the previously unrecognized role of archaeal Hsp60 in stabilizing membrane fluidity.

We conducted a similar series of experiments using DMPG LUV, mimicking the lipid composition of eukaryotic membranes. DMPG LUV revealed a rapid decrease in anisotropy with increasing temperature. Oligomeric Hsp60β attenuated this reduction, demonstrating its protective role in maintaining membrane integrity. Monomeric Hsp60β, in contrast, showed no significant impact on DMPG lipids (Fig 3p). Additionally, we examined Hsp60β’s ability to associate with DMPG by incrementally increasing its concentration. Our observations revealed that Hsp60β could indeed associate with DMPG LUVs revealed from increased DPH anisotropy (Fig 3q), with a calculated dissociation constant (K_d_) of 119.33±6.22 nM (Fig 3r). The higher K_d_ value compared to archaeosome interactions is consistent with the idea that archaeosomes are specific lipid membranes, while DMPG represents a more non-specific interaction for Hsp60β. To this end, the present findings contribute to our understanding of Hsp60β’s versatile functions in maintaining membrane integrity under varying conditions.

### Hsp60**α** forms hetero-oligomeric complex with **β** subunit and Hsp60**αβ** complex receives assistance from Hsp14

The group II chaperonin of *Sulfolobus* consists of three subunits: α, β, and γ. These subunits come together to create a diverse range of unique complexes with distinct functions. In our study, we also sought to delineate the characteristics of the Hsp60α subunit and its interactions with Hsp60β. FRET between Alexa-488 tagged α subunit and Alexa-532 tagged β subunit revealed a progressive increase in energy transfer as the concentration of β subunit increased (Fig 4a). The titration experiments gave us a robust binding affinity (Fig 4b), quantified by a K_d_ value of 167.37±1.7 nM (supplementary equation 5), indicative of a strong interaction between the two subunits.

**Figure 4:**
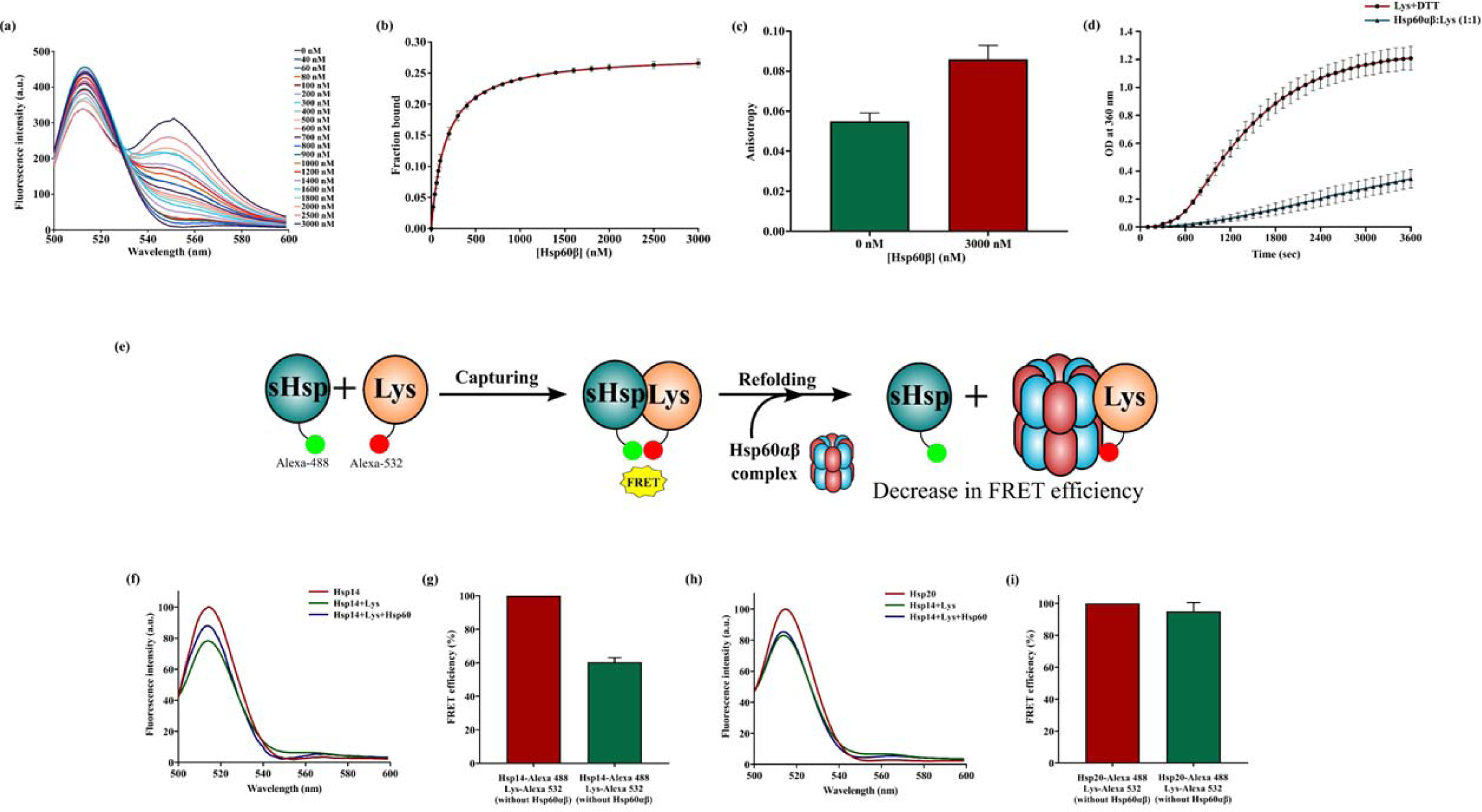
Diverse aspects of the Hsp60αβ complex. (a) The interaction between Alexa-488 tagged Hsp60α and Alexa-532 tagged Hsp60β was investigated through FRET, demonstrating their association. (b) Binding affinity between Hsp60α and Hsp60β was quantified using equation 5, yielding a K_d_ value of 167.37±1.7 nM. (c) In the absence of Hsp60β, the α subunit displayed low anisotropy. However, in the presence of Hsp60β at saturating concentrations, a significant increase in anisotropy was observed. (d) The Hsp60αβ complex exhibited the ability to protect lysozyme from DTT-induced damage, highlighting its functional importance. (e) Substrate transfer from small heat shock protein to Hsp60αβ complex was studied through a FRET based approach. The sHsps were tagged with Alexa-488 and lysozyme was tagged with Alexa-532. In the capturing state denatured lysozyme was allowed to get captured by sHsps resulting in FRET. However, in the refolding stage when Hsp60αβ complex was added substrate transfer resulted in decreased FRET efficiency. (f) Alexa-488 labelled Hsp14 was initially excited at 490 nm, and the resulting emission (red trace) was recorded within the 500 nm to 600 nm range. Subsequently, it was mixed with Alexa-532 labelled lysozyme, and the emission spectra were recorded (green trace). The presence of quenching in the spectra signifies the occurrence of FRET. Notably, upon the introduction of unlabelled Hsp60αβ (blue trace), FRET cancellation was observed, indicating the release of the substrate from Hsp14. (g) To determine the extent of FRET dissolution, the recorded data were first subjected to normalization and baseline subtraction. This process revealed that the FRET efficiency had decreased to 60%. (h) Alexa-488 labelled Hsp20 was excited at 490 nm, and the emission spectra (red trace) were recorded between 500 nm and 600 nm. Subsequently, it was mixed with Alexa-532 labelled lysozyme, resulting in quenching (green trace). Addition of unlabelled Hsp60αβ (blue trace) did not have any effect, indicating no substrate release from Hsp20. (i) After the recorded data were normalized and baseline subtraction was performed, the FRET efficiency had reduced to only 95%. Error bar represents SEM obtained from three or more sets of replicates.

We also assessed the anisotropy change of the hetero-oligomeric αβ complex. Without Hsp60β, the α subunit displayed a low anisotropy value. However, when Hsp60β was present at a saturating concentration, the anisotropy value significantly increased (Fig 4c). This notable change suggests the successful formation of a hetero-oligomeric complex in the presence of Hsp60β, underlining the importance of this interaction in the biological context. To examine the hetero-oligomeric αβ complex’s activity, we assessed its ability to protect a substrate from aggregation under stressful conditions. Lysozyme, prone to rapid aggregation in the presence of DTT, displayed reduced aggregation when the Hsp60αβ complex was present, highlighting its ability to safeguard substrates from environmental stressors (Fig 4d). Not only does it protect a substrate it can also refold a substrate back to its functional conformation. We tested its ability to refold heat inactivated calf intestinal phosphatase and observed almost 63% recovery in its activity (supplementary figure S2).

In our previous study, we showed involvement of small heat shock protein in substrate transfer to Hsp60β homo-oligomer [54]. We also wanted to identify the small heat shock protein (sHsp) responsible for mediating substrate transfer to the Hsp60αβ complex. To investigate this, we selected lysozyme as our substrate of interest and adopted a FRET-based approach. In this methodology, we labeled the small heat shock proteins, Hsp14 or Hsp20, with the donor fluorophore Alexa-488, while the lysozyme substrate was labeled with the acceptor fluorophore Alexa-532 (Fig 4e). We treated the lysozyme with the reducing agent DTT for 1 hour to induce substrate denaturation. Our study encompassed two distinct stages: the capturing and refolding stages. During the capturing stage, we monitored the potential capture of the substrate by the small heat shock proteins (Fig 4e). Such capture events would be accompanied by FRET, leading to the quenching of donor fluorescence. In the refolding stage, we examined whether substrate transfer occurred upon introducing the Hsp60αβ complex into the reaction. If transfer occurred, it would manifest as a reduction in the FRET efficiency (Fig 4e).

In our experiment, our initial focus was on confirming the occurrence of FRET between Alexa-488-labelled small heat shock protein (sHsp) and Alexa-532-labelled lysozyme. This was established by observing a decrease in donor fluorescence (Fig 4f). When Hsp14 was present in capturing stage, we observed a decrease in the donor fluorescence intensity confirming substrate capturing. Later in the experiment, when we introduced Hsp60αβ, we noticed the donor fluorescence increased (Fig 4f). This increase indicated that the FRET between Alexa-488-labeled Hsp14 and Alexa-532-labeled lysozyme was disrupted due to the release of the substrate from Hsp14. The FRET efficiency decreased by 40% (Fig 4g).

However, the same pattern did not manifest when Hsp20 was utilized. In the case of Hsp20, during the capturing stage, there was a noticeable decrease in emission spectra of Alexa-532 labeled Hsp20 (Fig 4h), implying successful substrate capture. However, when Hsp60αβ was subsequently added, the anticipated cancellation of FRET and subsequent increase in donor fluorescence did not occur (Fig 4h). The decrease in FRET efficiency was only 5% (Fig 4i). These results strongly suggest that Hsp14 plays a role in transferring the substrate to the Hsp60αβ complex for refolding.

### ATPase activity of Hsp60**β** and Hsp60**αβ** complex exhibits nested cooperativity and is a mosaic of group I and group II chaperonin

There lies a nested allosteric cooperativity in the ATPase activity of Hsp60β and Hsp60αβ complex. Various models have been formulated to elucidate the mechanism of cooperativity in ligand binding by oligomeric proteins. The Monod-Wyman-Changeux (MWC) model [55] and the Koshland-Nemethy-Filmer (KNF) model [56] are prominent examples. In both models, cooperativity arises from ligand-induced conformational changes, which can be either concerted (MWC), sequential (KNF), or combined. The concept of nested allostery was initially introduced to explain certain linkage phenomena in hemoglobin [57]. Nested allostery is particularly valuable when applied to large oligomeric proteins with a hierarchical structure. The hierarchical structure within these proteins implies the potential existence of a corresponding hierarchy in allosteric interactions. Previously proposed nested allosteric models include those solely based on the MWC formalism [58, 59] and others where KNF-type allosteric transitions are nested within MWC transitions [60, 61]. In the context of this study, we have observed nested cooperativity in ATP hydrolysis by Hsp60β and Hsp60αβ concerning ATP. In this scenario, positive cooperativity within the ring driven by MWC-type transitions is nested within a negative cooperativity between the rings driven by KNF model.

#### Positive Cooperativity in ATP Hydrolysis driven by MWC model

Positive cooperativity in ATP hydrolysis by Hsp60β (Fig 5a) and Hsp60αβ (Fig 5b) was investigated by measuring the initial rates of ATP hydrolysis over a range of ATP concentrations spanning from 0 to 1 mM. The data could not be accurately fitted into a single hill equation (Fig 5b inset). To address this, we employed a supplementary equations 1 and 2 to better model the observed behavior. Two distinct allosteric transitions were observed: one occurring at relatively low ATP concentrations (≤ 200 µM) and the other at higher ATP concentrations (Fig 5a-b). The midpoints of these transitions were found to be approximately 74.48±3.86 µM and 175.59±3.87 µM, respectively for Hsp60β and for Hsp60αβ complex the midpoints of the transitions were 54.07±0.113 µM and 746.35±45.1 µM respectively. The Hill coefficient for the allosteric transition at low ATP concentrations (≤ 200 µM) was determined to be 2.027±0.0781 and 2.665±0.189 for Hsp60β and Hsp60αβ respectively. This value aligns with previously reported values for group I and group II chaperonin respectively [62–64] and corresponds to the Hill coefficient of the first ring. Notably, the Hill coefficient for the second ring was found to be 9.69±1.11 and 7.872±0.114 for Hsp60β and Hsp60αβ respectively, indicating the presence of intra-ring positive cooperativity in the second ring and a strong negative cooperativity between the rings. This values closely approaches the upper limit of the Hill coefficient, which is equivalent to the total number of ATP-binding sites within a ring which is 9 for Hsp60β and 8 for Hsp60αβ. The observed positive cooperativity in ATP hydrolysis by Hsp60β at low ATP concentrations reflects the transition from the TT state to the TR state. This positive cooperativity pertains to the first level of allostery within the rings.

**Figure 5:**
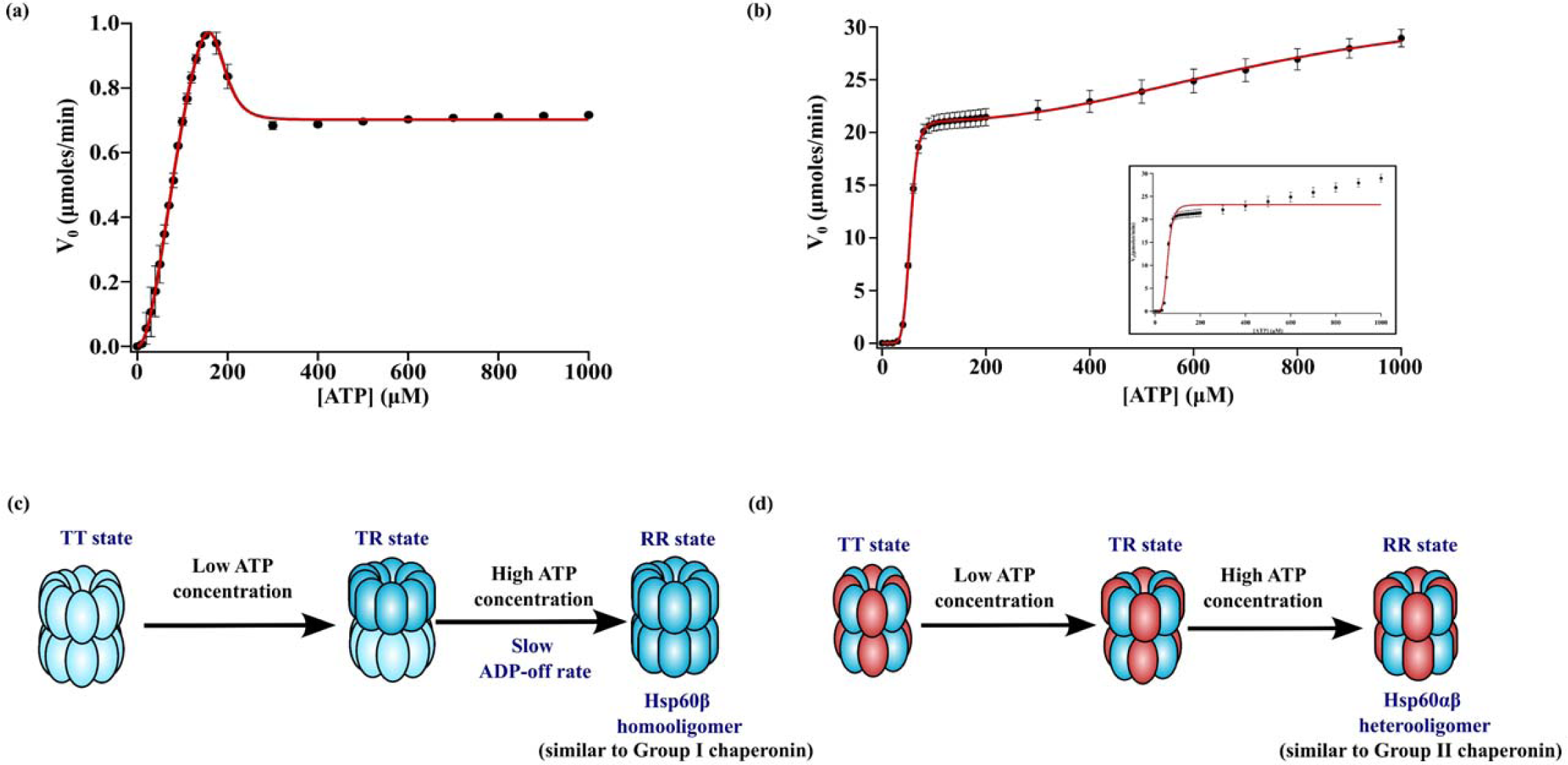
Mosaic nature in ATPase activity of Hsp60β and Hsp60αβ complex. The initial velocity of ATP hydrolysis by the Hsp60β homooligomer (a) and Hsp60αβ heterooligomer (b) was measured at various ATP concentrations. The experimental data were fitted to supplementary equations 1 and 2, with the Hsp60β oligomer concentration maintained at 90 nM. All reactions were conducted at a temperature of 70 °C, following the procedures outlined in the supplementary information. ATPase activity of Hsp60αβ could not be fitted into a single Hill equation (b inset). Notably, the resulting curve for Hsp60β revealed a slower ADP off-rate as ATP concentrations increased. However, Hsp60αβ complex did nit exhibit a slow ADP-off rate. (c and d) Scheme representing the transition of Hsp60β and Hsp60αβ particle from TT state to RR state through TR state. Within the rings there is a strong positive cooperativity driven by the MWC model and between the rings there is a negative cooperativity driven by the KNF model. At low concentration the particle transitions from TT state to TR state and at higher ATP concentration in transitions to RR state. Hsp60β had a slow ADP-off rate at higher concentration of ATP similar to group I chaperonin whereas Hsp60αβ did not exhibit such trait resembling group II chaperonin. sError bar represents SEM obtained from three or more sets of replicates.

#### Negative Cooperativity between the Two Rings driven by KNF model

The second level of allostery pertains to interactions between the two rings of the Hsp60β particle. The negative cooperativity between the two rings in ATP hydrolysis was assessed, yielding a Hill coefficient of 0.0044 and 0.00015 for Hsp60β and Hsp60αβ respectively, indicating a strong degree of negative cooperativity. Also, the negative cooperativity between rings, seems to be much stronger in Hsp60αβ than in Hsp60β. This negative cooperativity substantially impacts the overall positive cooperativity observed within the rings and the presence of two rings contributes to the biphasic nature of the ATPase activity curve.

Our findings propose a nested allosteric model to elucidate the cooperativity observed in ATP hydrolysis by Hsp60β and Hsp60αβ. This model comprises two distinct levels of allostery within the chaperonins: one operating within each ring and the other governing interactions between the two rings. These two levels of allostery align with the hierarchical structure of Hsp60β and Hsp60αβ. At the first level of allostery, each ring exists in an equilibrium between the T and R states, which follows the MWC representation [55]. The transition of the first ring from the T state to the R state, signifying the transformation of the Hsp60β and Hsp60αβ particle from the TT state to the TR state (Fig 5c-d), is evident through the positive cooperativity observed in ATP binding and hydrolysis by Hsp60β, primarily at relatively low ATP concentrations (≤ 200 µM). Conversely, the transition of the second ring from the T state to the R state, corresponding to the shift of the chaperonin particles from the TR state to the RR state (Fig 5c-d), becomes apparent when extending the range of ATP concentrations to 1 mM. This involves a sequential, KNF-type transition that progresses from the TT state to the asymmetrical TR state and culminates in the RR state. This nested allosteric model captures the intricate cooperative mechanisms within Hsp60β and Hsp60αβ, offering a more comprehensive understanding of its functionality.

Another important thing apart from the allostery is that the ATPase activity curve for Hsp60β displays a distinctive peak at approximately 200 μM ATP (Fig 5a). Notably, this curve shares qualitative similarities with the biphasic curves observed in the wild-type GroEL14 [65]. In the case of GroEL, the decline in ATPase activity at higher ATP concentrations has been attributed to the slow off-rate of ADP (Fig 5c). This phenomenon was subsequently found absent in the E257A GroEL mutant, which has a rapid off-rate for ADP [66]. Consequently, it is reasonable to hypothesize that the diminished ATPase activity observed with Hsp60β at higher ATP concentrations may also be attributed to a slow off-rate of ADP.

However, when we investigated the ATPase activity of the Hsp60αβ hetero-oligomeric complex, we made a separate observation. Our assessment of the ATPase activity of the Hsp60αβ hetero-oligomeric complex revealed nested cooperativity in the ATPase activity, with an ADP dissociation rate that was not slow at high ATP concentration (Fig 5b and d). This particular characteristic is typically associated with the ATPase activity observed in the TRiC/CCT hetero-oligomeric complex found in eukaryotic organisms [64]. This observation is unique as a single chaperonin molecule displays behavior pertaining to both group I and group II chaperonin depending on the nature of the complex formed. The varied ATPase activity in the Hsp60β and Hsp60αβ complex suggests a mosaic nature that may encompass features typical of both group I and group II chaperonins. The mosaic ATPase activity may reflect a functional diversity that allows these chaperones to adapt to different cellular environments and client proteins. This differential behavior of the Hsp60β homooligomeric and Hsp60αβ hetero-oligomeric complexes highlights that, archaea is indeed a mosaic of bacteria and eukaryotes. This mosaic nature is not only confined to their phylogenetic evolution alone but also extends to their cellular machinery, further highlighting the uniqueness of this domain of life.

A pivotal, unanswered question is whether ATP-induced conformational changes genuinely transpire in *Sulfolobus*, where the concentration of ATP is notably high, often within the millimolar range. An intriguing possibility arises: the concentration of ATP within the cavity of chaperonin may not reach equilibrium with the rest of the cell due to the sequestration of the chaperonin’s ATP binding sites, rendering them potentially inaccessible when unfolded proteins are bound. This arrangement of cooperativity in ATP hydrolysis by Hsp60β and Hsp60αβ might have evolved to facilitate responsiveness to small changes in the local ATP concentration during its reaction cycle. These changes could, in turn, trigger switches between different functional states, thus promoting the binding or release of protein substrates.

## Conclusion

Our study delved into the comprehensive structural and functional characteristics of Hsp60 (Hsp60β and Hsp60αβ) derived from the thermoacidophilic crenarchaeon *S. acidocaldarius* (Fig 6). This chaperone protein belongs to the group II chaperonin class and assumes its characteristic double-ring structure. The formation of these oligomers is dependent on the presence of ATP. These oligomeric Hsp60 structures (Hsp60β and Hsp60αβ) play a pivotal role in shielding proteins from aggregation under stress conditions by encapsulating them within their protective cage. Additionally, this versatile chaperonin can assist in refolding substrate proteins that have unfolded due to stress, helping them attain their native conformation. The folding mechanism probably follows an iterative annealing mechanism, in which Hsp60 captures the substrate based on hydrophobic patches present on the inner surface of its cage and folds them through charge driven repulsive forces. By doing so, Hsp60 stabilizes the folded form of the substrate, which generally possesses lower energy than the unfolded state, facilitating the substrate’s transition to its native structure. Hsp60 exhibits a unique form of nested cooperativity in its ATPase activity. This phenomenon features KNF-type cooperativity within the nested framework of the MWC model. The first ring displays positive cooperativity within itself at low ATP concentrations, while the second ring exhibits positive cooperativity at higher ATP concentrations. However, between the two rings, there is a strong negative cooperativity. Furthermore, a significant aspect of Hsp60 is its altered ATP hydrolysis behavior at higher ATP concentrations. Unlike the behavior of Hsp60β, which resembles the GroEL complex in bacteria with a slow ADP off rate, Hsp60αβ does not exhibit this behavior, aligning more with the TRiC/CCT complex from eukaryotes. This unique difference in ATP hydrolysis by archaeal thermosome highlights its distinctiveness. Depending on whether it forms a homo-oligomer or a hetero-oligomer, the ATPase activity of Hsp60 resembles either bacterial or eukaryotic chaperonins, showcasing the mosaic nature of archaea. The primary role of the Group I chaperonin GroEL is to refold unfolded substrates during stress. Interestingly, the Hsp60β homo-oligomer seems to emulate the functionality of the bacterial Group I chaperonin, particularly in response to elevated temperatures. Its primary function is to preserve protein homeostasis, a critical task during thermal stress. Additionally, Hsp60β exhibits the capability to safeguard membranes from thermal stress. In contrast, the functions of the eukaryotic CCT complex are diverse. Various subunits within the complex assume distinct roles, potentially facilitating the folding of different substrates and modulating interactions within and between rings [67]. This versatility underscores the complexity of the CCT complex. The Hsp60αβ complex resembles the CCT complex due to its two distinct subunits and it is reported to be formed under native growth conditions [21]. It might be possible that its role is to recognize different substrates and maintain protein homeostasis under normal growth condition. But this speculation requires further investigation. Consequently, we propose that the Hsp60 complex from *S. acidocaldarius* acts as a functional mosaic, combining attributes of both Group I and Group II chaperonins. This characteristic highlights the unique nature of archaeal domains.

**Figure 6:**
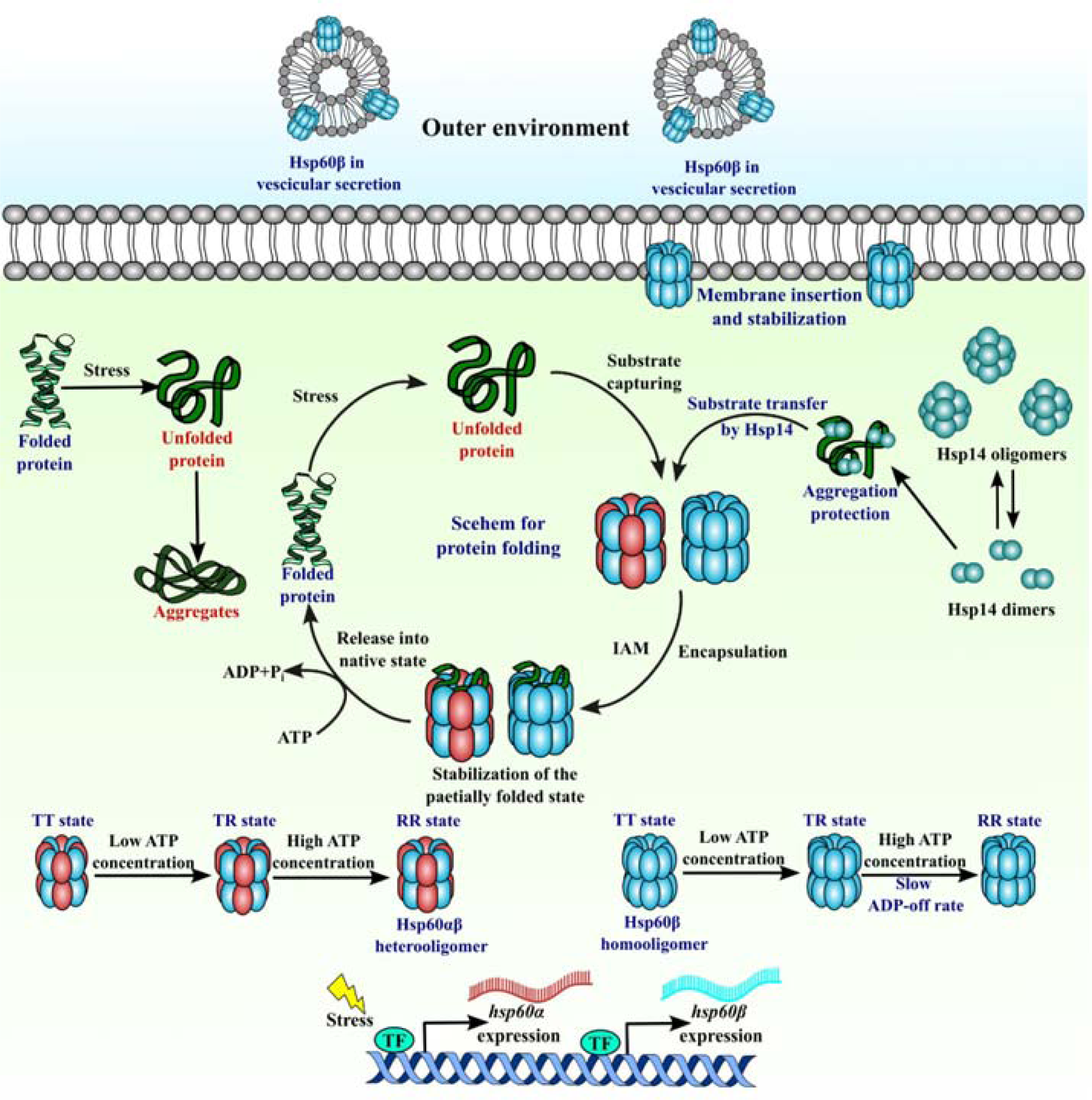
Model depicting heat shock response machinery in *S. acidocaldarius*. In response to stress, specific transcription factors play a pivotal role in orchestrating the positive upregulation of heat shock genes, such as hsp60α and hsp60β. This upregulation leads to elevated mRNA production, resulting in an increased concentration of Hsp60 within the cells. Hsp60, in turn, assembles into either homo-oligomeric or hetero-oligomeric structures in the presence of ATP, exhibiting a remarkable nested cooperativity in its ATPase activity. Hsp60’s functional role extends to its ability to discern unfolded substrate proteins based on their charge distribution, encompassing neutral and negative charges located along the inner surface of its protective cage. Once recognized, Hsp60 effectively stabilizes the folded state of the substrate, contributing to the restoration of proper protein structure. It also receives assistance from Hsp14. Furthermore, Hsp60 plays a crucial role in safeguarding cellular membranes against damage induced by stress. Its ability to protect membranes underscores its significance as a multifaceted molecular guardian, ensuring both protein integrity and membrane stability under adverse environmental conditions.

## Materials and methods

The supplementary information provides all the details of the materials used and methods.

## Acknowledgement

The authors would like to acknowledge the intramural funding from Bose Institute, India. KB was supported by a Senior Research Fellowship from Bose Institute, Kolkata. The authors would like to thank Dr. Utpal Bakshi for his support regarding analysis of the charge distribution of the *Sulfolobus* proteome. The authors would also like to thank Prof. Ajit Bikram Dutta for allowing the usage of fluorometer.

