## Supplementary information for "Functional diversity in the Hsp60 of *Sulfolobus acidocaldarius*: mosaic of Group I and Group II chaperonin"

**Materials and methods**

***Cloning and protein purification***

The genes *hsp60α* and *hsp60β* were cloned into the plasmids pETDuet-1 and pET28a, respectively. For hsp60α, the restriction enzymes SacI and SalI were used, while for hsp60β, NcoI and XhoI were employed. These genes were placed under the control of the T7 promoter and tagged with a C-terminal histidine tag. This resulted in the creation of two plasmids, namely pAG304 (containing *hsp60α*) and pAG305 (containing *hsp60β*). Subsequently, these plasmids were transformed into C43 (DE3)-RIL cells. Protein purification was conducted using a 1-liter culture. When the optical density at 600 nm (O.D.600) reached approximately 0.6, the cells were induced with 500 µM IPTG (Isopropyl β-D-1-thiogalactopyranoside) and incubated at 16ºC overnight with stirring to minimize the formation of inclusion bodies.

The following day, the cells were harvested and resuspended in a lysis buffer composed of 50 mM Tris (pH 8.0), 200 mM KCl, and 10% glycerol. Lysozyme and 1 mM PMSF (Phenylmethylsulfonyl fluoride) were added to the suspension, which was then kept on ice for 30 minutes. Subsequently, the cells were lysed through sonication, followed by centrifugation at 20,000 rpm for 30 minutes. The resulting supernatant was subjected to a heat treatment at 75ºC for 30 minutes and centrifuged again at 20,000 rpm for 30 minutes. The supernatant was then incubated with a Ni-NTA (Nickel Nitrilotriacetic Acid) column for 1 hour. The flow-through fraction was collected, and the column was subsequently washed successively with 10 mM and 20 mM imidazole solutions. The target protein was finally eluted using a 100 mM imidazole solution and subsequently dialyzed against the lysis buffer.

***Oligomerization of Hsp60β***

Hsp60β was subjected to heat treatment at 80ºC for 10 minutes in the presence of 5 mM ATP and 25 mM MgCl_2_. Following this treatment, the protein was loaded onto a Sephacryl S-300 column, which had been pre-equilibrated with 1 column volume (1CV) of lysis buffer. The protein was then eluted from the column using the lysis buffer at a flow rate of 0.5ml/min. The fractions collected during elution were subsequently subjected to SDS-PAGE (Sodium Dodecyl Sulfate-Polyacrylamide Gel Electrophoresis) to confirm the presence and purity of the protein.

***Intrinsic tryptophan fluorescence measurement***

Hsp60β was examined using intrinsic tryptophan fluorescence in the presence of 5 mM ATP and 25 mM MgCl_2_ under heated conditions. To conduct this experiment, the samples were excited at a wavelength of 280 nm, and the fluorescence emission spectra were measured at various time intervals, spanning from 300 nm to 400 nm. Data points were collected with a wavelength increment of 1 nm. This experimental setup allows for the monitoring of changes in tryptophan fluorescence over time, providing insights into the structural and conformational alterations of Hsp60β in response to ATP and MgCl_2_ under elevated temperature conditions.

***ATPase activity assay***

To determine the ATPase activity, 10 μg of Hsp60β was incubated in a buffer consisting of 50 mM HEPES (pH 7.5), 50 mM potassium acetate, 5 mM magnesium acetate, 1 mM DTT, and 5% glycerol at 70°C for 5 minutes. After the heat treatment, ATP was mixed with the reaction in varying concentration from 10 µM to 1000 µM and the reaction was het treated at 70 ºC for another 30 minutes. The released phosphate was measured using malachite green coloured reagent. The absorbance was measured at 620 nm. The data for the whole range of ATP was fitted into the following equation.

equation 1

$$V_{0}=\frac{V_{max(1)}+V_{max(2)}({\frac{[S]}{K_{2}})}^{m}}{1+({\frac{K_{1}}{[S]})}^{m}+({\frac{[S]}{K_{2}})}^{m}}$$

where V_0_ is the observed initial rate of ATP hydrolysis, [S] is the substrate (ATP) concentration, V_max(1)_ and V_max(2)_ are the respective maximal initial rates of ATP hydrolysis by a single ring and by both rings of thermosome, n and m are the respective Hill coefficients for ATP binding to the first and second rings, and K_1_ and K_2_ are the respective apparent binding constants of ATP for the first and second rings.

To determine the allosteric constants L_1_ and L_2_, the data was fitted into the following equation.

equation 2

$$V_{0}=\frac{{0.5V}_{max(1)}L_{1}({\frac{[S]}{K_{R}})(1+\frac{\left[ S \right]}{K_{R}})}^{N-1}+V_{max(2)}L_{1}L_{2}({\frac{[S]}{K_{R}})(1+\frac{\left[ S \right]}{K_{R}})}^{2N-1}}{1+L_{1}(1+{\frac{\left[ S \right]}{K_{R}})}^{N}+L_{1}L_{2}(1+{\frac{[S]}{K_{R}})}^{2N}}$$

Analysis of the second level of allostery between the two rings of chaperonin was performed by the following equation

equation 3

$$n_{50}=\frac{1}{1+({{L_{1}}/{L_{2})}}^{1/2}}$$

The Hill coefficient for two symmetric dimers and for interaction between two rings at 50% saturation is given by n_50_. The Hill coefficient depends on intrinsic allosteric constants L_1_ and L_2_.

***Tagging of recombinant proteins***

In the process of conjugating ALEXA-488 and ALEXA-532 fluorescent probes with purified recombinant proteins, a meticulous protocol, in accordance with the manufacturer's guidelines from Sigma-Aldrich, was meticulously followed. Initially, a solution of purified recombinant proteins was prepared, and subsequently, a sodium bicarbonate solution was added to create an optimal environment. The fluorescent probes' stock solutions were then carefully prepared in anhydrous dimethyl sulfoxide (DMSO). Protein labeling involved a slow addition of the fluorescent probes into the protein solution, followed by gentle stirring for 2 hours. Separation of labeled protein and unbound probe was achieved through gel filtration using a PD-10 column, and the collected fraction underwent a 24-hour dialysis against sodium phosphate buffer to eliminate excess probe and DMSO. The concentration and degree of labeling were assessed by measuring absorbance and fluorescence, with the Fluorophore-to-Protein (F/P) ratio indicating optimal labeling. A final confirmation of successful protein labeling was obtained through SDS/PAGE and scanning the gel using a Typhoon scanner. This comprehensive protocol ensures the efficient and optimized labelling of recombinant proteins, making them suitable for various assays and experiments requiring fluorescent detection.

***Fluorescence resonance energy transfer (FRET)***

To assess the interaction between Hsp60α and Hsp60β, we employed Fluorescence Resonance Energy Transfer (FRET) as our method of choice. Specifically, we used Alexa 488-tagged Hsp60α at a concentration of 500 nM, which was subjected to titration with Alexa 532-tagged Hsp60β at a temperature of 60°C. The resulting reaction mixture was excited at 490 nm, and emission spectra were meticulously recorded within the 500 nm to 600 nm range using a JASCO f-8500 spectrophotometer sourced from JASCO International Co., Ltd. in Japan. The emission spectra for both the donor and acceptor fluorophores were separated and analyzed using Igor Pro 6.0 software by WaveMetrics in the USA. The observed decrease in donor fluorescence intensity was plotted as a function of Hsp60β concentration, representing the fraction of Hsp60α bound to Hsp60β. Subsequently, this dataset was fitted to a mathematical equation for further analysis and interpretation.

equation 4

$$f_{b}=\frac{a \times[Hsp60\beta]}{[Hsp60\beta]+K_{d}}$$

where a is a constant, f_b_ is the bound fraction of Hsp60β, and K_d_ is the dissociation constant.

***Dynamic light scattering***

To assess the oligomeric state of Hsp60β at various temperatures, Dynamic Light Scattering (DLS) experiments were conducted using a Zetasizer Nano S instrument from Malvern Instruments in Malvern, UK. A sample concentration of 10 μM was prepared and passed through a 0.22-μm filter to remove any particulate matter.

Measurements were performed at two different temperatures in absence and presence of ATP, specifically at 70°C. Data were acquired as the average of 60 scans. The hydrodynamic radius (R_H_) of the Hsp60β particles was determined using the Stokes–Einstein equation, which relates the Brownian motion of particles to their hydrodynamic radius, allowing for an estimation of their size based on their diffusion behavior in solution.

equation 5

$$R_{H}=\frac{k_{B}T}{6\pi\eta D}$$

where k_B_ is Boltzmann’s constant, T is absolute temperature, η is the medium’s viscosity, and D is the translational diffusion coefficient of the particles.

***ANS study***

Fluorescence of ANS (20 µM) bound to 10 µM Hsp60 was measured at 60ºC at different time interval. In the fluorescence spectroscopy experiment, an excitation wavelength of 390 nm was employed. Fluorescence emission spectra were recorded within the wavelength range of 400–600 nm. Data were collected with a resolution of 1 nm for each data point. The slits for both excitation and emission were set at 5 nm. The reported spectra represent the average of three individual scans.

***Preparation of archaeosome***

To create archaeosomes for testing the interaction of Hsp60 with membrane lipids, *S. acidocaldarius* cells were cultured aerobically in Brock medium at a pH of 3 and a temperature of 76°C. Archaeal membranes were isolated using the Bligh-Dyer method. Initially, a solvent for extraction was prepared by mixing chloroform with methanol in a 2:1 ratio (for archaea). Then, 3.75 ml of this extraction solvent was added to 1 ml of the cell suspension. The mixture was vigorously vortexed for 10 minutes. Subsequently, 1.25 ml of chloroform was added and vortexed for an additional 1 minute. Finally, 1.25 ml of water (H_2_O) was added to the mixture, which was then centrifuged at 1000 rpm for 10 minutes. The organic layer, containing the lipids, was collected. The chloroform was removed from the collected organic layer using a Turbovap LV concentration evaporator (Biotage, Sweden) under N_2_ gas pressure at 65°C. The resulting membrane was resuspended in a 50 mM Sodium phosphate buffer at pH 8.0, preheated to 60°C. The resuspended membrane was rapidly frozen in liquid nitrogen (N_2_) and then thawed in a water bath at 60°C. Archaeosomes were prepared from the isolated membranes by subjecting the solution to sonication in a soniprep 150 sonicator for 5 minutes (with 10 cycles of 30 seconds each). The concentration of lipids was quantified using the Stewart assay. For *S. acidocaldarius* (Saci) lipids, the lipid concentration was determined to be 0.3 mg/ml.

***Preparation of DMPG LUV***

To prepare Large Unilamellar Vesicles (LUVs), DMPG (Dimyristoylphosphatidylglycerol) lipid was purchased from Avanti Polar Lipids Inc. in Alabama, USA. A total of 5 mg of the DMPG lipid was accurately weighed and then dissolved in 300 µl of chloroform, which was sourced from Merck KGaA in Darmstadt, Germany. The lipid solution was dried by directing a stream of nitrogen gas over it. Subsequently, the lipid was lyophilized overnight to form a lipid film. This process removes the solvent and leaves behind a lipid film. The next day, the lipid film was hydrated by adding 1 ml of 50 mM Tris buffer with a pH of 8.0 and containing 100 mM NaCl. The mixture was vortexed for 30 minutes. To achieve non-uniform sized vesicles, the hydrated lipid solution underwent five freeze-thaw cycles. This involved alternating between freezing the solution in liquid nitrogen and thawing it in lukewarm water. The vesicles were then extruded through a 100-nm polycarbonate membrane filter obtained from Avanti Polar Lipids Inc. (Alabama, USA). A mini extruder from Avanti Polar Lipids in Alabaster, Alabama, was used for this process. The extrusion setup was configured to produce LUVs with a diameter of approximately 90-95 nm. The suspension obtained after extrusion was ready for use in subsequent experiments. These LUVs are essential for studying various biological and chemical processes, especially those involving interactions with lipids.

***Archaeosome and DMPG stabilization assay***

A 10 mM solution of DPH (1,6-Diphenyl-1,3,5-hexatriene) (Sigma) was prepared in DMSO. It was then added to a 100-fold diluted sample of archaeosome or 50 µg/mL DMPG solution with the final DPH concentration being 3 μM. This was followed by incubation at 16 °C for 10 min. Anisotropy of DPH in presence of oligomeric or monomeric Hsp60β or without any protein was then measured in a JASCO fluorimeter (Jasco, Japan) at various temperatures 30°C, 40°C, 50°C, 60°C and 70°C for archaeosome and 20°C, 25°C, 30°C, 35°C and 40°C. Anisotropy percentage was calculated by considering anisotropy at 30°C or 20°C as 100% and plotted against temperature for each of the three conditions using Sigma plot 12.0.

***Titration of archaeosome or DMPG***

After labelling the archaeosome or LUV, it was subjected to titration with an increasing concentration of Hsp60. The fluorophore 1,6-Diphenyl-1,3,5-hexatriene (DPH) was used, and its excitation occurred at 350 nm. The resulting anisotropy was measured within the emission wavelength range of 400 nm to 500 nm.

Changes in the anisotropy (r) were quantified using the following equation, which helps assess the rotational mobility of the labeled molecules:

equation 6

$$r=\frac{I_{ǁ}-I_{\perp}}{I_{ǁ}+2I_{\perp}}$$

where I_||_ is intensity of detected light when the excitation and emission polarization are parallel and I_⊥_ is intensity of detected light when the excitation and emission polarization is perpendicular. The fraction bound was calculated from the measured anisotropy using the following equation:

equation 7

$$f_{b}= \frac{r-r_{free}}{r_{bound}-r_{free}}$$

where, f_b_ represents the bound fraction of labeled molecules, which indicates the proportion of labeled molecules that have interacted or bound with Hsp60, stands for the measured anisotropy at a specific concentration of Hsp60. This value reflects the degree of rotational mobility or freedom of the labeled molecules in the presence of Hsp60, r_free_ represents the anisotropy observed in the absence of Hsp60. It serves as a baseline measurement, indicating the rotational mobility of labeled molecules when no binding or interaction with Hsp60 has occurred, and r_bound_ signifies the anisotropy after saturation, which indicates the maximum restriction of rotational mobility of labeled molecules when they are fully bound to Hsp60. To analyze the data, the fraction bound is calculated for various concentrations of Hsp60, representing the proportion of labeled molecules interacting with Hsp60 at each concentration. These values are then plotted against the increasing concentration of Hsp60. Typically, this data is fitted using a hyperbolic binding equation. This is a mathematical model often used to describe the binding behavior of ligands to a receptor. In this context, it helps describe how the interaction between Hsp60 and labeled vesicles results in changes in rotational mobility, and the equation provides a means to quantify this binding behaviour.

***Aggregation protection***

10 µM lysozyme was mixed with 20 mM DTT. DTT is a thiol based reducing agent which reduces disulfide bond and leads aggregation of protein. Light scattering at 360 nm was measured. Aggregation protection activity of either oligomeric or monomeric form Hsp60β was measured in ratio 0.5:1 and 1:1 with respect to lysozyme.

***Calculating the charge distribution of Sulfolobus proteome***

To calculate acidic/basic nature of the proteomes, the following set of four rules was fed into the designed algorithm:

**Rule 1:** The condition to confer acidic or basic nature to proteome is as follows: i) if the isoelectric point (pI) of a protein is <6; the protein is acidic in nature ii) if the pI of a protein is >=6 to <8; the protein is neutral in nature; and iii) if pI of a protein is >=8; the protein is basic in nature.

**Rule 2:** The calculation of pI is based on different pKa sets and the Henderson–Hasselbach equation as implemented in the following models: Solomon (1), Lehninger (2), EMBOSS (3), Dawson (4), Toseland (5), Sillero (6), Thurlkill (7), Rodwell (8), DTASelect (9), Nozaki (10), Grimsley (11), Bjellqvist (12), ProMoST (13) and IPC (14).

**Rule 3:** To compute consensus pI values, the average of all pI calculations from each model is considered. However, if a protein's consensus pI deviates by more than 5% from the average value of all models, the pI of that protein is excluded from the final computation (marked as red in the output matrix).

**Rule 4:** The resulting output provides a frequency table indicating the distribution of acidic, basic, and neutral nature across the proteomes of the four organisms.

**Table S1: Strains, Primers and plasmids used in the study**

| **Bacterial strains, primers and plasmids** | **Relevant characteristics** | **Source of reference** |
| --- | --- | --- |
| **Bacterial strains** | | |
| *E. coli* XL1Blue | *recA1 endA1 gyrA96 thi-1 hsdR17 supE44 relA1 lac* [*F´ proAB lacIq ZΔM15 Tn10* (*Tet^r^*)]  Mesophilic organism which grows optimally at 37 °C | Stratagene |
| *E. coli* BL21(DE3)-RIL | *B F-ompT hsdS*(*rB−mB−*) *dcm + Tet^r^ E. coli gal λ* (*DE3*) *endA Hte* [*argU ileY leuW Cam^r^*]  Mesophilic organism which grows optimally at 37 °C | Stratagene |
| **Archaeal strain** | | |
| *Sulfolobus* *acidocaldarius*  DSM 639 | Hyperthermo-acidophilic crenarchaeon which grows optimally at 80°C and pH 2–3 | DSMZ |
| **Primers** | | |
| 88 | 5’-GGGGGCCATGGGCATGGAGCTTCACATATG-3’ Forward primer for *saci_1665* containing a NcoI restriction site (underlined) | [1] |
| 89 | 5’-GGGGGCTCGAGTTCGATCTTAATTGCTGATAC-3’ Reverse primer for *saci_1665* containing a XhoI restriction site (underlined) | [1] |
| 74 | 5’-AAAAACCATGGGCATGTCAAGAAGATATAG-3’ Forward primer for *saci_0922* containing a NcoI restriction site (underlined) | [2] |
| 75 | 5’-AAAAACTCGAGCTCAACTTTAATTTCAGTACC-3’ Reverse primer for *saci_0922* containing a XhoI restriction site (underlined) | [2] |
| 321 | 5’-GGGGGCCATGGGCATGTCAGCTACAGCTACAGTTG-3’ Forward primer for *saci_0666* containing a NcoI restriction site (underlined) | [1] |
| 322 | 5’-GGGGGCTCGAGGTCTTCTTCTTTACC-3’ Reverse primer for *saci_0666* containing a XhoI restriction site (underlined) | [1] |
| 317 | 5’-GGGGGGAGCTCGATGGCTGGTCCAGTTC -3’ Forward primer for *saci_1401* containing a SacI restriction site (underlined) | The present study |
| 318 | 5’-GGGGGGTCGACTCACTCTAATGAAGG-3’ Reverse primer for *saci_1401* containing a SalI restriction site (underlined) | The present study |
| **Plasmids** | | |
| pET28a | Kan^r^, Bacterial expression vector with T7lac promoter containing replicon ColE1 (pBR322) and one MCS. | Novagen |
| pAG15 | Kan^r^, pET28a carrying C terminal His-tagged *saci_0922* (sHSP) using restriction sites NcoI-XhoI | [2] |
| pAG17 | Kan^r^, pET28a carrying C terminal His-tagged *saci_1665* (sHSP) using restriction sites NcoI-XhoI | [1] |
| pAG305 | Kan^r^, pET28a carrying C terminal His-tagged *saci_0666* (Hsp60 β subunit) using restriction sites NcoI-XhoI | [1] |
| pAG304 | Ampr, pETDuet1 carrying C terminal His-tagged *saci_1401* (Hsp60α subunit) in MCS1 using restriction sites SacI-SalI | The present study |

**Results:**

***The ATPase activity of Hsp60β increases in presence of substrate***

The specific recognition and binding to denatured proteins or nonnative intermediates represent critical steps in chaperone function [3]. In order to assess whether non-native proteins could indeed induce ATP hydrolysis, we conducted experiments to measure ATPase activity in the presence of varying concentrations of two distinct denatured substrates: lysozyme and MDH. Our findings indicated that ATPase activity exhibited a proportional increase with rising concentrations of both lysozyme (ranging from 10 to 70 µM) (Fig S1a) and MDH (from 1 to 7 µM) (Fig S1b). Notably, for lysozyme, the enhancement in ATPase activity was particularly pronounced and did not appear to reach a saturation point even at the highest tested concentration of 70 µM. Conversely, in the case of MDH, we observed that initially, at lower MDH concentrations, the ATPase activity remained relatively low. However, as the MDH concentration increased, the ATPase activity demonstrated a significant surge and eventually seemed to plateau at approximately 7 µM MDH concentration. From these results, we can reasonably conclude that the binding of non-native lysozyme and MDH effectively activated ATP hydrolysis, surpassing the basal steady rate of ATPase activity. This demonstrates the importance of these chaperone proteins in recognizing and interacting with denatured or nonnative substrates to facilitate essential cellular processes.

We conducted experiments to investigate the influence and metal ion dependency on the ATPase activity of Hsp60β (Fig S1c). Our observations revealed a notable dependency on metal ions for ATPase activity. In the absence of any metal ions, we observed minimal to negligible ATPase activity. However, when metal ions were introduced, we found significant variations in ATPase activity levels. The highest ATPase activity was observed in the presence of Mn^2+^, followed closely by Co^2+^. This finding aligns with an earlier report that documented the activation of ATPase activity in Cpn from *Pyrococcus furiosus* by either Co^2+^ or Mn^2+^ [4]. Interestingly, ATPase activity remained relatively consistent in the presence of Ca^2+^ and Mg^2+^. Notably, a Group II chaperonin from *Thermoplasma acidophilum* was shown to display superior ATPase activity and refolding efficiency when exposed to Mn^2+^ compared to Mg^2+^ [5]. Conversely, the presence of Ni^2+^ resulted in the lowest observed ATPase activity for Hsp60β. These findings underscore the sensitivity of ATPase activity in Hsp60β to different metal ions and highlight the importance of specific metal ions, such as Mn^2+^ and Co^2+^, in modulating the chaperone's functional capabilities.

We investigated the impact of temperature on the ATPase activity of Hsp60β. In the absence of a denatured substrate, we observed a relatively low basal ATPase activity of the protein (Fig S1d). However, the presence of denatured, non-native lysozyme led to a notable increase in ATPase activity. The ATPase activity exhibited a distinct temperature dependency, with a peak observed at 80°C (Fig S1d). Beyond this point, as the temperature continued to rise, the ATPase activity gradually decreased. At 95°C, we recorded the lowest ATPase activity, indicating a significant structural perturbation of the protein, which consequently led to a reduction in its activity. This suggests a critical temperature threshold beyond which the protein experiences substantial structural alterations that impair its functionality.

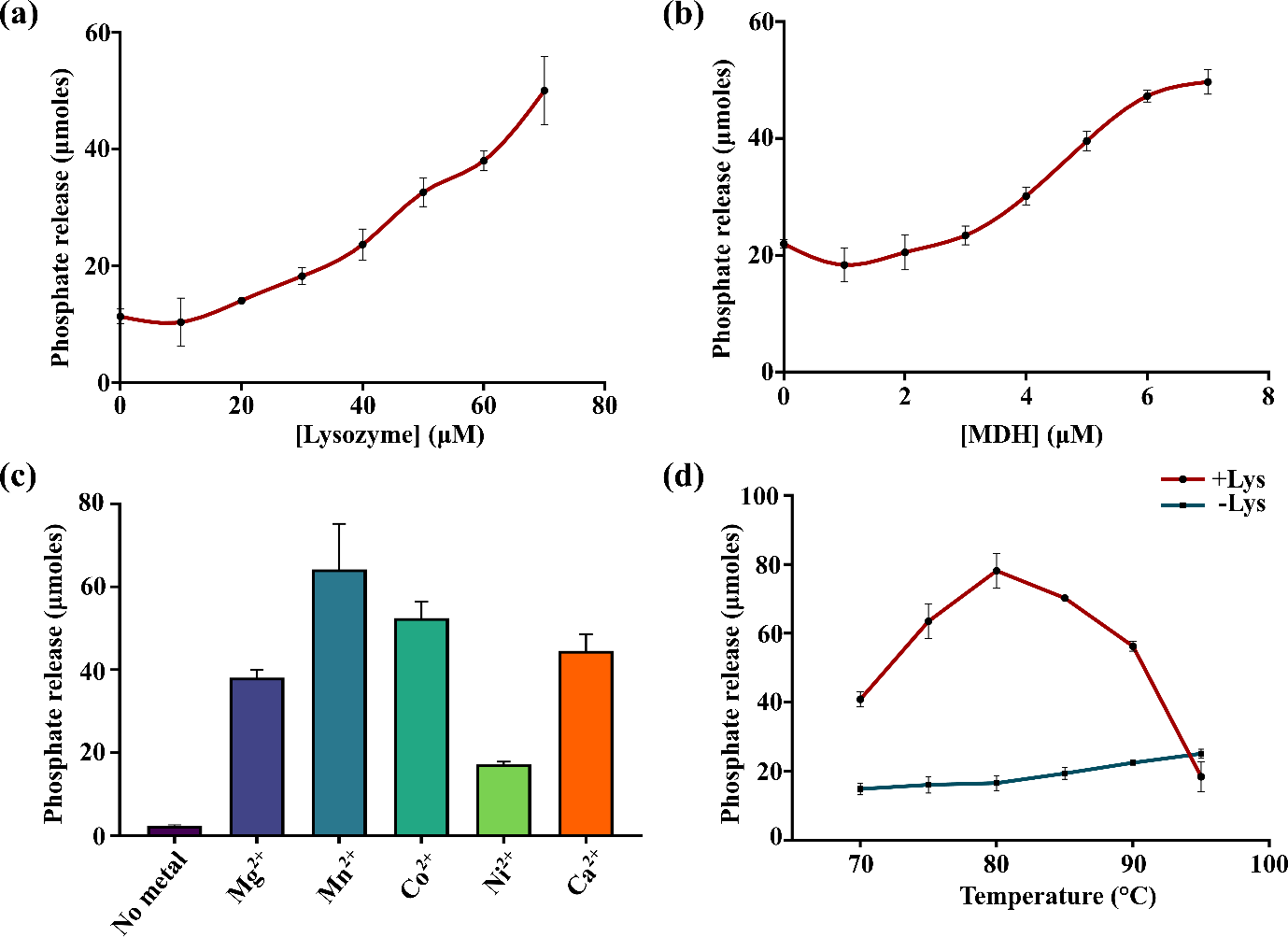

**Figure S1.: The ATPase activity of Hsp60β exhibits an upward trend as substrate concentration increases.** The ATPase activity of Hsp60β was assessed in the presence of escalating concentrations of lysozyme (a) and MDH (b). In both cases, the release of phosphate increased proportionally with higher substrate concentrations. (c) The assessment of ATPase activity was extended to various divalent metal ions, with the most pronounced activity observed in the presence of Mn^2+^. (d) Further evaluation involved measuring ATPase activity at different temperatures, both with and without the presence of lysozyme as a substrate. In the presence of lysozyme, the highest ATPase activity was recorded at 80 ºC, while in its absence, the ATPase activity remained nominal.

***Hsp60αβ can refold a substrate back to its native state***

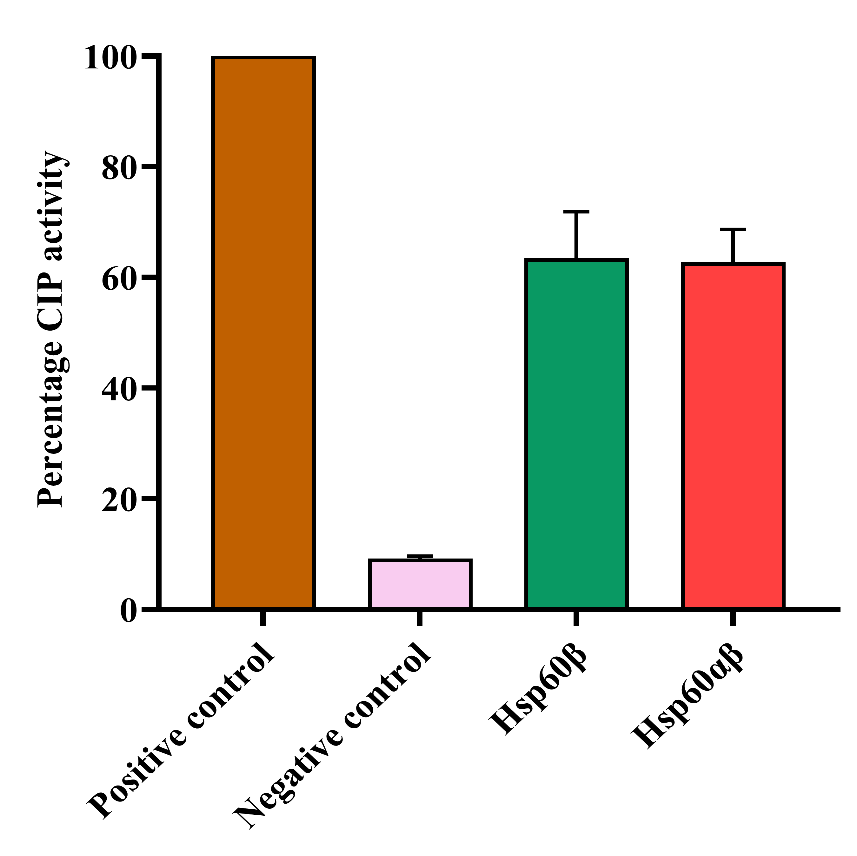
We assessed the refolding capacity of the Hsp60αβ complex using calf intestinal phosphatase as the substrate. Calf intestinal phosphatase (CIP) is an alkaline phosphatase derived from the gastrointestinal mucosal barrier of calves. The enzymatic activity of CIP was quantified by employing para-nitrophenyl phosphate (pNPP) as a substrate. The active CIP catalyzed the cleavage of phosphate from pNPP, generating para-nitrophenol (pNP), which exhibits a yellow color. The absorption of pNP at 420 nm allowed us to measure CIP activity by recording the optical density at that wavelength. Additionally, CIP is sensitive to heat, and a brief heat treatment at 60 °C for only 10 minutes resulted in a reduction of the protein's helical content, leading to protein unfolding [6]. We observed that in the negative control where CIP was heat treated without the addition of Hsp60β or Hsp60αβ complex the activity reduced to 9%. But in the presence of either Hsp60β or Hsp60αβ complex the activity was restored to nearly 63% suggesting that both can refold a heat inactivated substrate to its native conformation.

**Figure S2.: Hsp60αβ can refold heat inactivated CIP*.*** Positive control represents the sample without heat treatment, and its activity is considered to be 100%. Negative control represents a heat-treated sample without any Hsp60 complex. In presence of either Hsp60β or Hsp60αβ complex the activity was restored. Error bar denotes SEM from three or more data sets.
